# Cold and lonely: low microbial abundance and no core microbiota in mosquito populations across Greenland

**DOI:** 10.1101/2024.08.27.609970

**Authors:** Diana Rojas Guerrero, Tomas Roslin, Otso T. Ovaskainen, Michał Kolasa, Viktor Gårdman, Karol H. Nowak, Monika Prus-Frankowska, Mateusz Buczek, Anna Michalik, Piotr Łukasik

**Author notes:** Correspondence: Diana Rojas Guerrero, and Piotr Łukasik. Authors’ email addresses: DRG,; TR,; OTO,; MK,; VG,; KHN, MPF, MB,; AM,; PŁ,.

## Abstract

Microbial symbionts are key players in insect ecology and evolution, but microbiota vary widely across and within host species. The microbiota of insect clades such as high-latitude mosquitoes remain understudied, despite their abundance and potential economic and medical relevance. Here, we resolve temporal and spatial variation in the abundance and diversity of bacteria and fungi associated with *Ochlerotatus* mosquitoes in Greenland. To this aim, we apply a comprehensive workflow combining the simultaneous characterization of host, bacterial, and fungal marker regions, contamination filtering, as well as bacterial quantification and joint species distribution modelling.

We find that 573 mosquitoes collected in five regions of Greenland between 2009 and 2024 represented two species (*O. nigripes* and *O. impiger*). Their microbial communities were low-abundance, highly diverse, and dominated by environmentally versatile taxa, showing high individual-to-individual-rather than spatio-temporal variation, although taxa such as *Pseudomonas* and *Serratia* exhibited sex-specific associations. DNA from local vertebrates and potential vertebrate pathogens (e.g., *Bartonella*) was detected, likely reflecting blood meals. These findings suggest that Greenland *Ochlerotatus* mosquitoes do not rely on core microbiota, but engage in flexible associations with diverse, low-abundance microbial communities. They also point to mosquitoes as likely vectors of bacterial and fungal pathogens in the Arctic.

## 1. Introduction

Symbiotic microorganisms play crucial roles in the ecology and evolution of insects, often influencing host biology in multiple ways (Bhattacharyya et al., 2026; Douglas, 2022). While many insects rely on specialised, vertically-transmitted microbes for well-defined functions (i.e., nutrition, detoxification, protection against pathogens or parasitoids), others appear to thrive with sparse or absent host-adapted microbial communities (Hammer et al., 2017, 2019; Malacrinò, 2022), associating with transient microbes instead (Ravenscraft & Coon, 2026). As a consequence, microbial diversity and abundance can vary dramatically even within species (Lange et al., 2023; Malacrinò, 2022; Sanders et al., 2017), highlighting the dynamic and context-dependent nature of host-microbe relationships, and the need for their systematic study.

Mosquitoes (family Culicidae) are one of the insect clades whose microbiota have attracted substantial attention. While adults rely on carbohydrates (e.g. nectar and honeydew) as the primary energy source, the females of many species are also hematophagous, often feeding on blood of reptiles, birds and mammals, including humans (Barredo & DeGennaro, 2020; Corbet & Downe, 1966; M. Lee et al., 2025). Because many species can vector disease-causing pathogens such as bacteria, viruses, filarial worms, and protozoa (Hegde et al., 2015; Moonen et al., 2023; Petersen et al., 2015), they have been extensively studied as causes of disease outbreaks (e.g., malaria, dengue, Chagas) in tropical regions. As such, their interactions with microbes that could influence disease transmission became an important area of research.

Mosquitoes have been reported to harbour diverse microbial communities that, in some species, can influence various aspects of their biology, including development, immunity, and interactions with pathogens (Bai et al., 2019; Coon et al., 2014; Guégan et al., 2018; Hegde et al., 2018). For example, several species host the facultative heritable endosymbiont *Wolbachia*, which can influence the vectoring capacity, reproduction, and fitness of their mosquito host (Alomar et al., 2023; Hoffmann et al., 2024; Minwuyelet et al., 2023). Further, studies across mosquito microbiota have consistently identified members of the genera *Pseudomonas*, *Serratia*, *Enterobacter*, *Commamonas*, and many others, as well as fungi such as *Cladosporium* and *Aspergillus*, as common associates (Guégan et al., 2018; Hegde et al., 2024; Thongsripong et al., 2018). In contrast to insects that maintain stable, maternally or socially transmitted symbionts, microbial community composition in mosquitoes vary substantially among populations and life stages since many mosquito-associated microbes are thought to be environmentally acquired during larval development or adult feeding (Coon et al., 2016). As a result, mosquito microbiota may reflect local environmental conditions, microbial source pools, and ecological context. Nevertheless, much of the mosquito microbiome research has focused on a few model species and typically on laboratory populations, which has left the natural microbiome of the vast majority of >3,000 recognised mosquito species underexplored (Guégan et al., 2018; Hall, 2022). As a result, the ecological and evolutionary drivers of microbiota variation in natural mosquito populations remain poorly understood.

Arctic ecosystems, of which mosquitoes are often important components, provide a unique context for investigating the factors that shape insect-associated microbial communities. The Arctic is characterized by short growing seasons, low temperatures, strong seasonal fluctuations, and relatively simple ecological networks, all of which may influence the diversity and transmission of microbial associates. Additionally, limited species diversity facilitates the study of the relative importance of these drivers.

In the Arctic ecosystems of Greenland, two mosquito species are overwhelmingly abundant, *Ochlerotatus nigripes* (Fig. 1a,b) and *Ochlerotatus impiger* (both formerly *Aedes)* (Böcher, 2015; Reeves et al., 2013). They feed on the blood of a wide range of local mammals and birds, but also on plant nectar (Corbet & Downe, 1966), playing key roles in food webs and nutrient cycling.

**Figure 1.**
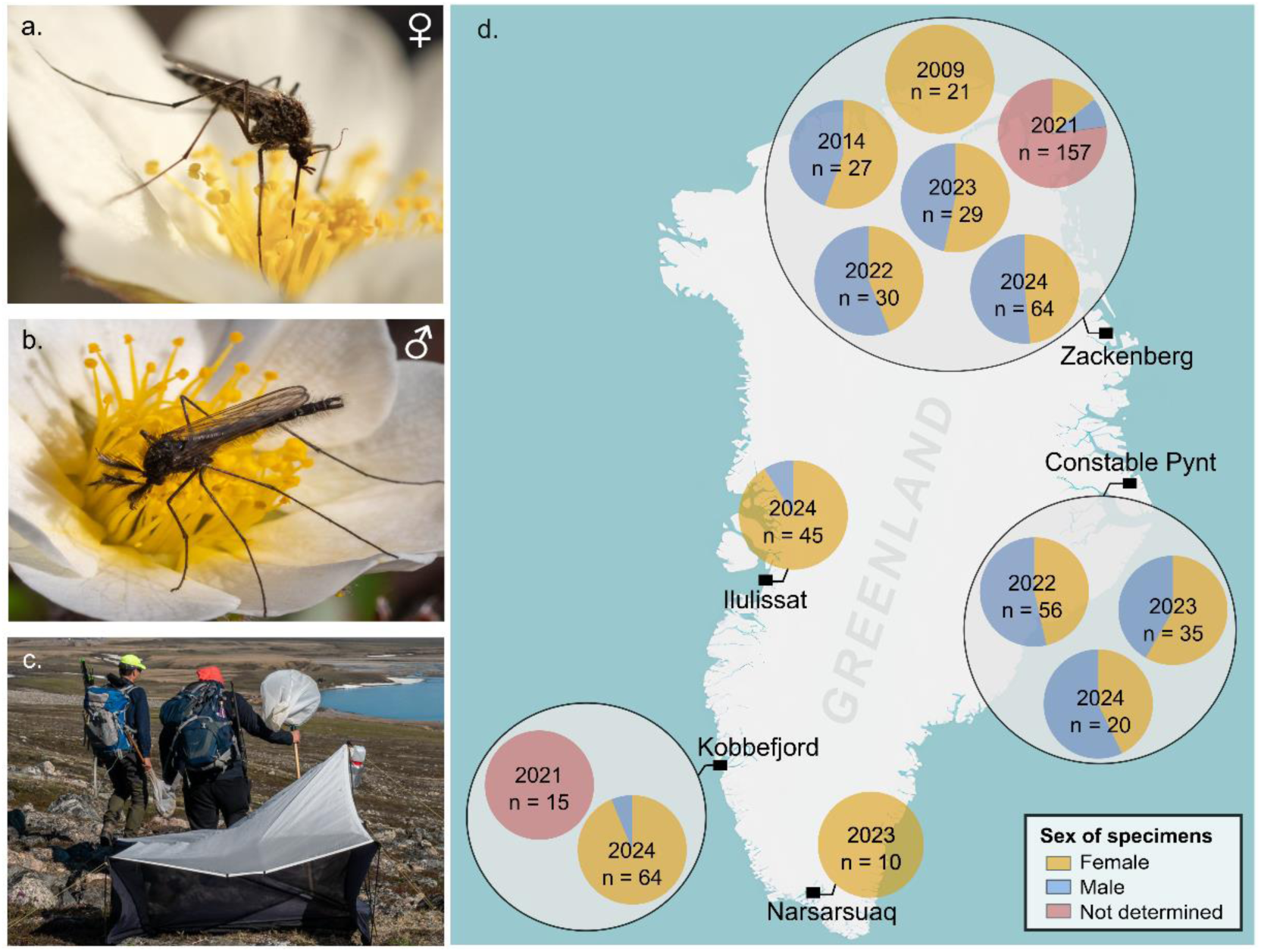
*Ochlerotatus nigripes* a) female and b) male, feeding on nectar of *Dryas* flowers in Zackenberg Valley, NE Greenland. c) The authors (TR & MK) in Zackenberg Valley, demonstrating the sampling tools: a Malaise trap and a sweep net. Photos: PŁ. d) Mosquito sampling sites, years, and sampling depth. Colors within pie charts indicate the proportion of successfully processed females, males, and individuals whose sex was not recorded, in each site/year combination.

Previous research on Greenland mosquito microbiome has focused primarily on their interactions with viruses. Although the overall evidence suggests that the vectorial capacity of *O. nigripes* and *O. impiger* is limited (Corbet, 1967; Müllerová et al., 2018; Schilling et al., 2025; Snyman et al., 2024), metagenomic work showed that mosquitoes display species-specific and regional viral community signatures. By contrast, host-bacterial and -fungal interactions have not been characterized for *Ochlerotatus* in Greenland, leaving open the question of whether such patterns can be observed in bacterial and fungal communities or whether they remain consistent across geographic regions.

In this study, we asked about the patterns and likely drivers of microbial abundance, diversity, and distribution across Greenland mosquitoes. By simultaneously amplifying marker genes to identify the mosquito species and characterize their associated fungal and bacterial communities, including bacterial abundance quantification and comparisons with other mosquito and other insect species, we profiled the microbiota of 573 individuals collected across five areas of the island between 2009 and 2024. We then used statistical modelling to disentangle the environmental and host-related factors underlying the variation of their microbial communities.

## 2. Materials and Methods

### 2.1. Sampling and specimen selection

Wild adult mosquitoes were collected by sweep netting or Malaise traps (Fig. 1c) in five areas of Greenland: Zackenberg Valley, Constable Pynt, Narsarsuaq, Kobbefjord, and Illulissat, between 2009 and 2024 (Fig. 1d, Table S1). Zackenberg Valley was surveyed repeatedly over multiple years, whereas additional locations were sampled during one or two seasons and incorporated to broaden the geographic coverage of the study. Specimens were preserved in 96% ethanol and stored at -20°C until processing.

Mosquitoes from Zackenberg 2021 and Kobbefjord 2021 were a part of large diverse insect collections (Buczek et al., 2024) processed without their prior identification or sexing; we later selected mosquito specimens based on the barcode sequence (see below). From other site/year combinations, we specifically selected and sexed mosquitoes based on morphology (Dahl, 2015). As a reference for microbial abundance comparisons, from the Zackenberg 2021 collection, we selected four other taxonomically diverse and well-represented insect species. As additional references for these comparisons, we used *Wolbachia*-infected and *Wolbachia*-free *Aedes aegypti* adult mosquitoes from a long-term laboratory culture at Jagiellonian University and overwintering *Culex pipiens* female mosquitoes collected in the village of Przybysławice near Miechów, South Poland (Table S1).

### 2.2. Amplicon library preparation and sequencing

Mosquitoes were processed individually, in five batches (Table S1), following the DNA extraction and library preparation protocol detailed recently (Buczek et al., 2024). In short, the mosquitoes were placed in 2 ml tubes with 200 μl lysis buffer, homogenised using ceramic beads, and then incubated at 55 °C for two hours. To one-fifth (40 μl) of the homogenate aliquot, we added 20,000 copies of a linearised plasmid carrying an artificial 16S rRNA amplification target (Ec5502) (Tourlousse et al., 2016), to enable subsequent quantification, and purified the DNA using SPRI beads (Rohland & Reich, 2012). For each batch of 88 samples, we used a positive extraction control and two blank extraction controls.

Amplicon libraries were prepared using a two-step PCR approach (Table S2). During the first step, we simultaneously amplified a 418 bp portion of insect cytochrome c oxidase I (COI) gene, the 16S rRNA V4 region (16S hereafter) for bacteria, and the Internal Transcribed Spacer region 2 (ITS2) for fungi, using a mixture of primers BF3-BR2 (Elbrecht et al., 2019), 515F-806R (Parada et al., 2016), and fITS1F-fITS2 respectively, with Illumina adapter tails. These primers were selected based on their broad coverage and extensive validation in metabarcoding surveys (Iwaszkiewicz-Eggebrecht et al., 2023; Thompson et al., 2017), and their compatibility in a multiplex reaction. The first-step PCR products were cleaned using SPRI beads and then used as templates for the second PCR reactions, which used primers that completed the adapters and added a unique combination of indexes to each sample. Two positive PCR controls and two blank PCR controls were included along with mosquito samples for PCR1, and an additional blank control was included for PCR2. Amplification success and amplicon size were verified by agarose gel electrophoresis after each PCR. All amplicon libraries were pooled together based on gel band intensity, cleaned with SPRI beads, and submitted for sequencing on either NovaSeq 6000 SPrime 500-cycle flow cell or NextSeq 2000 P2 600-cycle flow cell (Table S1).

### 2.3. Data analysis

Demultiplexed .fastq files for all samples were run through a bioinformatic pipeline (Buczek et al., 2024), described at https://github.com/Symbiosis-JU/Bioinformatic-pipelines. Initially, reads were split into bins corresponding to different targeted regions (COI, 16S rRNA, or ITS2) based on primer sequences at the beginning of each read. For each marker region, paired-end reads were quality-filtered and merged using PEAR (Zhang et al., 2014). Representative sequences were then dereplicated, denoised, taxonomically classified, and the resulting ASVs (Amplicon Sequencing Variant) were clustered into Operational Taxonomic Units (OTUs) using a 97% similarity threshold, using VSEARCH (Rognes et al., 2016), USEARCH (Edgar, 2010), and UNOISE (Antich et al., 2021). Taxonomy assignment was based on the MIDORI reference database version GB239 (Leray et al., 2022) for COI, the SILVA database v138 (Quast et al., 2012; Yilmaz et al., 2014) for 16S rRNA, and the UNITE database v10.0 (Abarenkov et al., 2024) for ITS2 sequences. For each region, the pipeline provided tables with two levels of classification: ASVs and OTUs (deposited at https://doi.org/10.6084/m9.figshare.c.8012695), both of which were subsequently used as input for a series of marker region-specific custom steps, as described below.

### 2.4. Mitochondrial data analysis

The ASV COI data were used to determine mosquito population structure across the sampled locations and to reconstruct potential biotic interactions. Initially, for all libraries with >100 *Ochlerotatus* COI reads remaining post-filtering, we selected the most abundant COI variant among ASVs taxonomically assigned to the genus *Ochlerotatus* as the specimen barcode. Libraries with zero or <100 COI mosquito reads were discarded from further analysis. The genetic distances and geographic distributions of haplotypes were visualized using PopART (Bandelt et al., 1999; Clement et al., 2002; Leigh & Bryant, 2015).

COI ASVs corresponding to taxa other than mosquitoes were separated and used to reconstruct the presence/absence of biological signal of associated species (i.e., putative sources of blood meal, or parasitic mites). We selected 10 reads as the minimal threshold for reporting non-human signal derived from blood meals as long as a) the confidence of the taxonomy assignment was >95%, b) corresponded to a species recorded in Greenland, and c) the signal was found in a library corresponding to a female mosquito, or a mosquito with sex not recorded. Regarding human detection, we only considered it reliable in three individual mosquitoes, one of which was captured and individually preserved after feeding on human host; all yielded ≥708 human COI reads, whereas none of the other samples had more than 10. Clean COI reads can be found in Supplementary File 3.

### 2.5. Microbial data analysis

We used the 16SrRNA OTU table for a custom reference-based decontamination process combined with template quantification (Łukasik et al., 2017; Nowak et al., 2025). We excluded any OTUs taxonomically assigned as mitochondria, chloroplasts, Eukaryota, Archaea, or chimeras. We then compared the maximum relative abundances of all OTUs across experimental samples and controls, assuming that true insect-associated microbes should be much better represented in insect samples than in blanks. Hence, any OTUs for which the maximum relative abundance attained in at least one experimental library did not exceed at least ten-fold the maximum relative abundance attained in any blank library were classified as a contaminant and excluded (Łukasik et al., 2017). Additionally, we filtered OTUs classified taxonomically as *Polaromonas* sp., *Brachybacterium* sp. or *Variovorax* sp., known reagent-derived contaminants in our laboratory. These reagent-derived bacteria comprised the large majority of reads in negative controls and those filtered from experimental samples (Fig. S1).

We estimated the absolute abundance of bacterial 16S rRNA gene copies in each processed insect individual based on the spike-in references (Tourlousse et al., 2016). Specifically, for all libraries, we multiplied the ratio of non-contaminant reads to spike-in reads by the number of spike-in copies added to the purified homogenate aliquot (20,000), and by 5, to account for using only one-fifth of the insect homogenate. To assess whether such calculated microbial load varied significantly across mosquito populations from various Greenland locations sampled over multiple years, and between sexes and mosquito species, we applied a generalised linear model (GLM) to log-transformed absolute 16S rRNA copy counts. Since we were primarily interested in the consistency of abundances across mosquito species, sexes, sites, and years, we included the main effects of these factors and their interactions. Models were fitted using the stats package (Venables & Ripley, 2002) in RStudio v4.4.1. Samples for mosquito individuals with undetermined sex were excluded from this analysis.

ITS2 OTU table was decontaminated as described for 16S rRNA data. As no ITS spike-ins were included during library preparation, we used the insect COI read number as a baseline and reference for comparing fungal amounts among individual insects (Płoszka et al., 2025). To this effect, for all libraries with fungal amplicon data, we computed ITS2/COI read number ratios as a means of comparing fungal load. Decontaminated bacterial and fungal OTU reads are provided in Supplementary File 3 (0.6084/m9.figshare.31934055).

OTU richness and abundance distribution were estimated for the full bacterial and microbial decontaminated datasets. Non-metric Multidimensional Scaling (NMDS) was performed using the ordinate function in the Phyloseq package (McMurdie & Holmes, 2013) with Bray-Curtis dissimilarity, four dimensions (k = 4), and up to 200 iterations per run (try = 200), allowing a maximum of 500 steps per iteration (maxit = 500), and up to 1000 random starts (trymax = 1000). Ordinations included only samples for which individual sex was determined. ComplexPheatmap (Gu, 2022; Gu et al., 2016) and ggplot2 (Wickham, 2016) were used for visualisation. Core microbiome analysis and UpSet plots were performed with mia (Borman et al., 2025) and microbiome (Lahti & Sudarshan, 2017).

### 2.6. Hierarchical Modelling of the Microbial Species Communities

To assess how host factors and environmental settings influence the composition of *Ochlerotatus* microbial communities, we used generalised linear mixed modelling of multivariate responses, applied through the joint species distribution model of Hierarchical Modelling of Species Communities (HMSC) (Ovaskainen, 2020; Ovaskainen et al., 2017). For bacteria, variation in 16S rRNA OTUs was quantified using spike-in-corrected reads across 400 mosquitoes estimated to contain at least 1000 copies of the gene, whereas fungal ITS2 OTUs were analysed using read counts across 364 libraries containing at least 100 reads.

Due to the zero-inflated nature of the data (i.e., the many absences of individual taxa, corresponding to counts of zero), we modelled microbial abundances through a hurdle approach: we first modelled presence-absences, and then abundance conditional on presence, fitting separate models for bacteria and fungi. In modelling presence-absences, we assumed a Bernoulli distribution and a probit link function; in modelling abundance conditional on presence, we log-transformed and scaled the response, assuming a normal distribution.

To measure the extent to which the mosquito microbiota composition is predictable, we used the species of mosquito, sex of individuals (excluding those for which sex was not recorded), and the interaction between sex and species as fixed effects. Because the sequencing depth of each individual (defined as decontaminated read counts) is likely to affect the detectability of microbial taxa, we included this covariate in the presence/absence models of both bacteria and fungi. For bacteria, variation in sequencing depth was adjusted for by relating the number of reads of each taxon to the reads from a known quantity of spike-in (see previous section), and no further correction for read counts was required. For fungi, where no spike-in benchmark was available, read counts were included as a covariate in the abundance model. As random effects, we included mosquito COI haplotype, sampling location, sampling year, and host individual to account for hierarchical structure and variation specific to each sampled mosquito.

We assumed the default prior distribution of HMSC (Ovaskainen, 2020; Ovaskainen et al., 2017) and fitted the model using the Markov Chain Monte Carlo (MCMC) approach implemented in the R-package Hmsc (Tikhonov et al., 2017). Model performance was evaluated in terms of explanatory and predictive power using AUC for presence-absence models and R^2^ for models on abundance conditional on presence.

To examine the importance of fixed and random effects in structuring microbiome variation, we applied a variance partitioning approach to the explained variation, quantifying the proportion of variance attributable to each factor and covariate. To examine the effects of sex and species, we evaluated the proportion of taxa that showed either a positive or negative response with at least 95% posterior probability. Residual patterns of co-variation among taxa, with at least 95% posterior probability, were examined across hierarchical levels (host individual, haplotype, location, and sampling year) to identify associations not explained by the measured covariates.

## 3. Results

Following multitarget amplicon sequencing for 609 adult mosquitoes, raw reads for each sample were split into amplicon datasets corresponding to COI, 16S rRNA, and ITS2 markers. Samples for which either COI or 16S rRNA amplification was unsuccessful were discarded. Ultimately, we retained amplicon libraries for a total of 573 mosquitoes, including 328 from Zackenberg, 111 from Constable Pynt, 10 from Narsarsuaq, 45 from Ilulissat, and 79 from Kobbefjord, sampled across different years (Fig. 1d).

### 3.1. Geographic distribution and biotic interaction patterns of Greenland mosquitoes

We first assessed the species diversity and distribution across sampled mosquito populations and years using mitochondrial COI amplicon sequencing data. After filtering out rare and noisy COI amplicon sequence variants (ASVs), we retained a total of 27,996,651 COI reads, 48,859 per library on average (range 104-370,010). Over 99% of these were assigned to mosquitoes, with reads comprising 569 ASVs clustered into 15 operational taxonomic units (OTUs), all belonging to the genus *Ochlerotatus*. Most ASVs were assigned to a single OTU, indicating that most individuals belonged to the same species-level lineage.

COI haplotype analysis based on the most abundant sequence variant in each mosquito individual, considered as its barcode, revealed 53 such variants across mosquito populations, including 35 assigned to *O. nigripes* and 18 to *O. impiger. O. nigripes* occurred at all sampled sites, whereas *O. impiger* was detected only in South and West Greenland. Across populations, each species was dominated by a single haplotype (Fig. 2a), with the dominant haplotypes differing by 10 of 418 bp (2.4%) between species. In each population, the dominant haplotype was frequently accompanied by several less common haplotypes, many of which were represented by only a single individual and were often separated from the dominant haplotype by a single nucleotide. Despite the presence of multiple haplotypes, within-species mitochondrial divergence was shallow, with low nucleotide diversity in both *O. nigripes* (*π* = 0.00064, *S* = 28, Tajima’s *D* = -2.42) and *O. impiger* (*π* = 0.00089, *S* = 18, Tajima’s *D* = -2.74), consistent with a structure dominated by one common haplotype and multiple rare derivatives.

**Figure 2.**
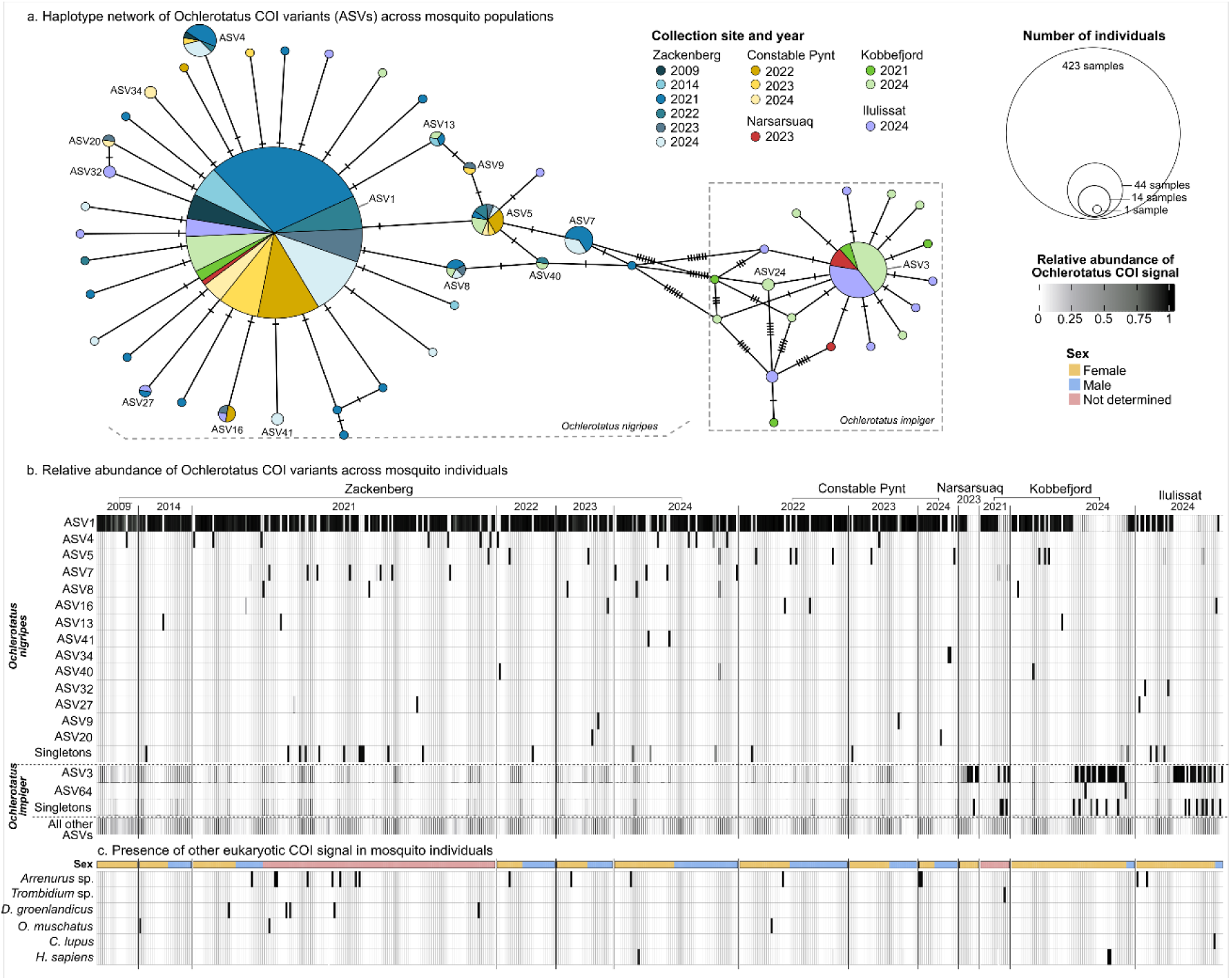
Mitochondrial diversity of *Ochlerotatus* mosquitoes and their biotic interactions. a) Minimum spanning network of *Ochlerotatus* barcodes. Circles represent unique haplotypes, scaled by the number of individuals, hatch marks indicate 1 bp differences, and colors denote sampling site and year. b) Relative abundance of the most common *Ochlerotatus* COI variants (ASVs) among *Ochlerotatus* mosquitoes from different sampling locations and years. Variants that were found in a single library were grouped and are presented jointly as ‘singletons’. c) The presence of reads representing non-mosquito animals in mosquito COI libraries. The sex of each individual is displayed by color in the top row. In panels b and c, columns represent individual mosquitoes.

Among the 0.41% of COI reads not corresponding to *Ochlerotatus* (Fig. 2c, Supplementary File 3: Tables a-b), the majority were assigned to the water mite genus *Arrenurus*, found across 16 mosquito individuals, while reads assigned to the mite genus *Trombidium* were found in one individual. Further, in 13 mosquitoes, we detected COI reads matching locally present mammals, likely traces of blood meals. This included the northern collared lemming (*Dicrostonyx groenlandicus)* detected in seven mosquito individuals, the muskox (*Ovibos moschatus*) detected in two individuals, the Greenland dog (*Canis lupus familiaris)* in one, and human (*Homo sapiens*) in three (Fig. S3). The mammalian signal was always found in female mosquitoes or in individuals whose sex was not determined, consistent with the expectation that only female mosquitoes feed on blood.

### 3.2. Low and variable microbial load across *Ochlerotatus* populations

For the 573 *Ochlerotatus* mosquitoes confidently classified into species, we obtained a total of 7,164,008 reads for the V4 region of the 16S rRNA gene, with an average of 12,503 reads per library (range 508 – 114,404). Most of these reads represented OTUs abundant in negative controls and corresponding to known reagent-derived contaminants, or alternatively, spike-in quantification plasmids (Fig. S1). Such reads were bioinformatically filtered using blank controls as references, and after this decontamination procedure, individual libraries retained an average of 1,250 reads (range 0 – 47,627)(Supplementary File 3: Table c). Unlike in studies that do not account for contamination, the low share and number of non-contaminant reads observed here were expected for low-biomass samples, in which contaminant-derived DNA can comprise a substantial but variable proportion of total reads (Williamson et al., 2025). The removal of contaminants resulted in markedly reduced read counts for *Ochlerotatus*, but not for most reference species processed at the same time using the same methods (Fig. 3, Fig. S1, Table S1). Next, based on spike-in to non-contaminant bacteria read number ratios, we estimated absolute 16S rRNA copy number in individual insects (i.e., absolute bacterial abundance). We inferred that *Ochlerotatus* mosquitoes harbour a median of 8,546 bacterial rRNA copies per individual (0–19,000,000) (Table 1). Across sampling sites and years, most populations showed similarly low bacterial loads, although a small subset of individuals exhibited substantially higher values (Fig. 3).

**Figure 3.**
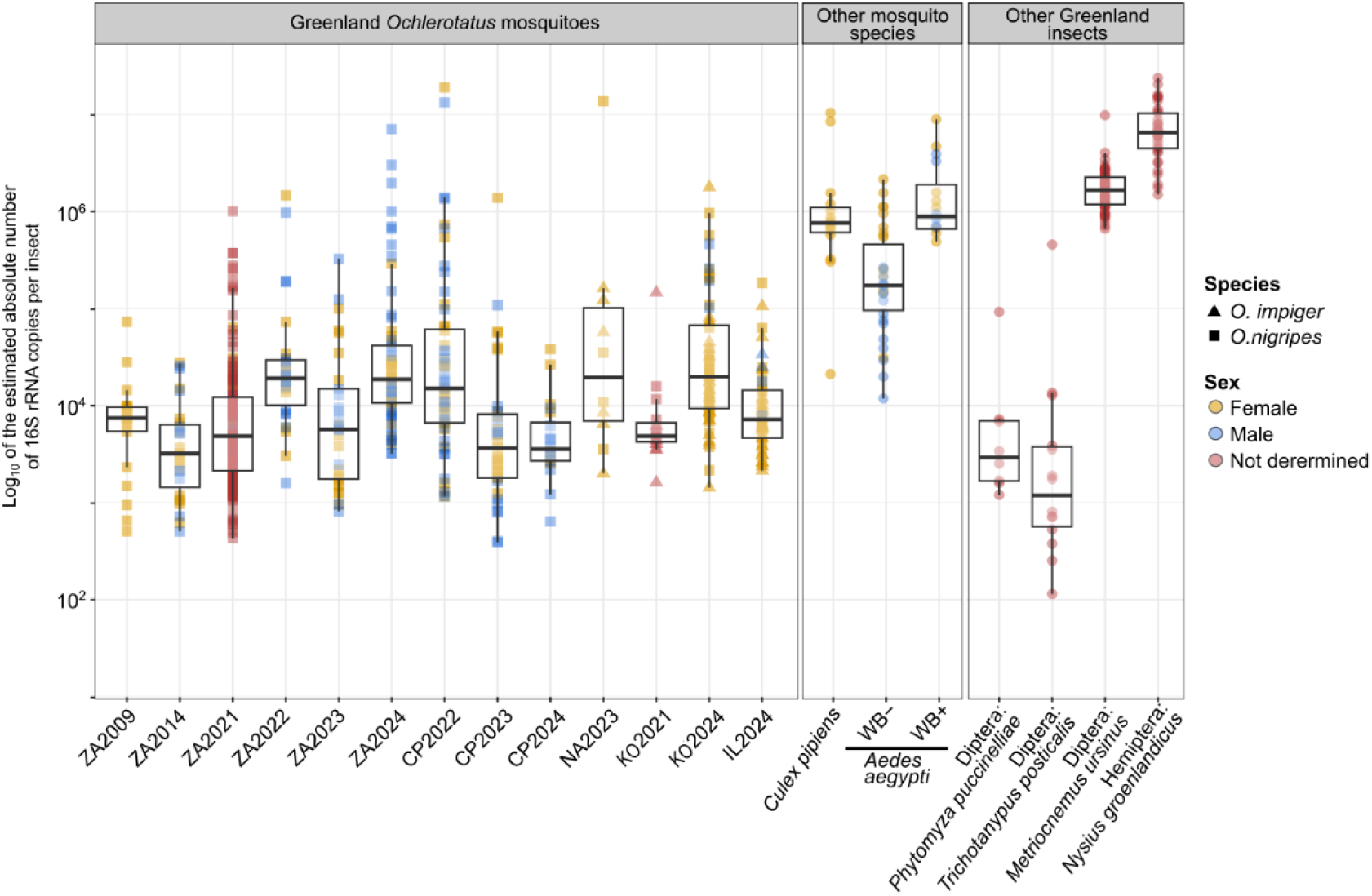
Absolute 16S rRNA abundance in *Ochlerotatus* mosquitoes from different sites and years, estimated based on quantification spike-in plasmids, was added to the homogenate and co-amplified alongside bacterial targets. Reference species include wild *Culex* mosquitoes from Poland, cultured *Aedes aegypti* mosquitoes from lines either infected (WB+) or non-infected (WB-) by *Wolbachia,* and four insect species collected in Zackenberg in 2021, distantly related to mosquitoes but of comparable body size.

**Table 1.** Bacterial 16S rRNA abundance across insect taxa: absolute abundances in individual insects estimated based on spike-in reads, and the abundance of non-contaminant 16S rRNA reads relative to COI reads.

| Species | n | Total 16S rRNA copies |  |  | Average<br>16S/COI ratio |
| --- | --- | --- | --- | --- | --- |
|  |  | Median | Mean | Range |  |
| *Ochlerotatus* spp. | 573 | 8,546 | 153,315 | 0 – 19,000,000 | 0.076 |
| *A. aegypti* WB- | 34 | 173,284 | 373,445 | 11,829 – 2,145,833 | 0.22 |
| *A. aegypti* WB+ | 16 | 892,208 | 1,926,381 | 489,900 – 8,926,860 | 1.7 |
| *C. pipiens* | 15 | 760,627 | 1,881,956 | 21,133 – 10,360,000 | 2.2 |
| *P. puccinelliae* | 8 | 2,972 | 14,977 | 1,202 – 92,260 | 0.03 |
| *T. posticalis* | 14 | 1,281 | 38,090 | 115 – 457,415 | 0.1 |
| *M. ursinus* | 56 | 1,668,170 | 2,418,833 | 663,060 -9,808,108 | 2.14 |
| *N. groenlandicus* | 37 | 6,517,603 | 7,861,297 | 1,496,034 – 23,834,590 | 13.57 |

To place these estimates in context, we compared them with other insects processed in the laboratory using identical extraction, amplification, and sequencing procedures (Fig. 3; Table 1). In other mosquitoes, including *Wolbachia*-infected (WB+) *Aedes aegypti* and *Culex pipiens* but also *Wolbachia*-free (WB-) *A. aegypti*, estimated bacterial abundances were between one and two orders of magnitude greater than in *Ochlerotatus* (Table 1, Fig. 3). Other Greenland insects like *Metriocnemus ursinus* and *Nysius groenlandicus* harboured median bacterial loads exceeding those of *Ochlerotatus* by more than two orders of magnitude. On the other hand, *Phytomyza puccinellia*e and *Trichotanypus posticalis* showed substantially lower abundances than *Ochlerotatus* (Fig. 3; Table 1).

A further, independent indication of low bacterial abundance in *Ochlerotatus* was provided by the ratio of bacterial 16S rRNA reads to COI reads, which were amplified simultaneously in the same multiplex PCR reactions, under identical conditions for different species. In *Ochlerotatus*, the average 16S/COI read number ratio was 0.076, whereas much higher ratios were observed in other mosquito species, including 0.22 in WB- *A. aegypti*, 1.7 in WB+ *Aedes aegypti*, and 2.2 in *C. pipiens.* The ratios varied from 0.03 to 13.57 in insects other than mosquitoes, in agreement with estimated 16S rRNA abundances (Table 1).

The GLM model revealed that total bacterial abundance varied in different ways across years in different sites (interaction site × year, X² = 29.50, p < 0.001) and between sexes across years (interaction sex × year, X² = 4.77, p = 0.029). No significant effects of sex, mosquito species, or their interaction were detected (Table S3).

### 3.3. High diversity and inconsistent bacterial community composition

Across the 573 *Ochlerotatus* libraries, we detected a total of 13,597 non-contaminant ASVs clustered into 4,328 OTUs, 945 genera, and 278 orders. Only 39 of 278 bacterial orders and 32 out of 945 genera occurred in more than 10% of individual mosquitoes. Among the orders that were both highly prevalent (detected in >50% of the libraries), and abundant (contributing >10% of rRNA copies in a library on average) were Burkholderiales (73.6% prevalence, 10% rRNA copies), Enterobacteriales (87.8%, 18.2%) and Pseudomonadales (63%, 10%) (Fig. 4a). The OTUs with the highest prevalence and abundance across all libraries corresponded to *Pseudomonas* (detected in 47.5% of libraries, 8% rRNA copies) and *Serratia* (41.2%, 5%). Some other OTUs, such as *Escherichia/Shigella, Enterobacter*, or *Janthinobacterium* were highly prevalent (52.4%, 33.2% and 28.4, respectively) but their relative abundance across libraries was low (4%, 3.3% and 2.2%, respectively) (Fig. 4b). These clades are known for their broad environmental distribution, including the ability to colonise diverse hosts, and four of the five have been listed among common contaminants in molecular biology reagents (Salter et al., 2014). However, strict contamination control applied in our study and the high ASV-level diversity observed in our data for these OTUs support a biological basis for these host-microbe associations (Fig. S1, S2). A distinct category comprises the bacterium *Bartonella,* which was detected in only seven mosquitoes (1.2% prevalence), but proved highly abundant in the samples where it was present, with an average of 526,772 16S rRNA copies per library (median = 351,795). Interestingly, in three of the mosquitoes carrying *Bartonella*, we detected COI signal of the northern lemming, a likely source of a bloodmeal and possibly the source of *Bartonella* infection (Fig. S3).

**Fig. 4.**
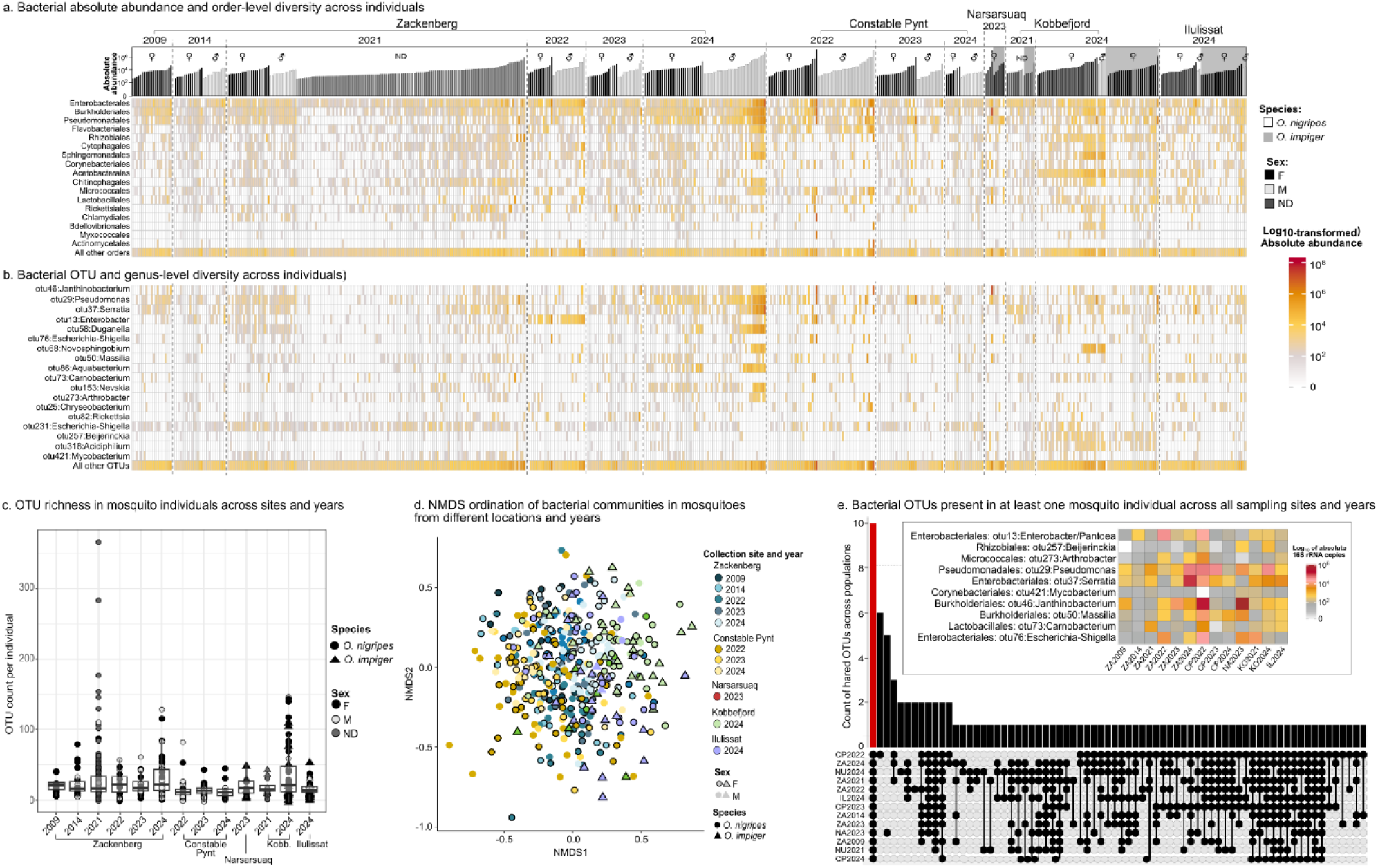
Bacterial diversity in mosquito individuals across sampling locations and years. a) OTU richness across sampled populations. Each point represents a number of OTUs in an individual mosquito, with the color indicating its sex and shape indicating species. b-c). Estimated absolute bacterial abundance (16S rRNA copy number) for all bacteria combined and for the most prevalent bacterial orders [b] and OTUs [c] across 573 mosquito individuals. Individuals are sorted by location, year, species, sex, and total bacterial abundance, in the same order for both panels. Only taxa with prevalence >35% across libraries are shown. d) NMDS ordination of community dissimilarity, based on Bray-Curtis distances, with females and males of the two mosquito species across sampled populations (site×year combinations). e) UpSet plot showing the presence/absence distribution of 100 most abundant bacterial OTUs across the dataset. The bars indicate the number of shared and unique OTUs, and the connected dots below the x-axis show which populations share each set. Population names are given as the first two letters of the site name and the year. The red bar highlights the number of taxa shared by all groups, and the inset heatmap displays the assigned taxonomy and the average absolute abundance of each OTU per sampled location/year.

On average, individual mosquitoes carried 26 OTUs (ranging from 0 to 366 OTUs in total) (Fig. 4c, Supplementary File 3: Tables c-d). The difference between the overall OTU count in the dataset and the OTU count across individual mosquitoes attests to remarkable variation in microbiome richness and diversity among individuals and populations. A non-metric multidimensional scaling (NDMS) based on Bray–Curtis dissimilarity matrix (stress value = 0.102) did not show clustering patterns by microbial composition across sampling locations or years (Fig. 4d). Further, based on the relative abundance threshold of 0.1% and a 75% prevalence, we were unable to detect a core microbiota within or among populations in any of the sampling years, locations, or sample-location combinations (Fig. S4). On the other hand, we detected a set of 10 OTUs (encompassing 7 orders and 10 genera) that were present in at least one individual in each collection (defined as site/year combination) (Fig. 4e).

### 3.4. Low abundance and inconsistent community composition in fungal microbiota

For 410 *Ochlerotatus* libraries with fungal data, we obtained 761,495 total ITS2 reads (1,894 per library on average, range 2-75,906), corresponding to 1,649 OTUs. Approximately 88% of these reads matched OTUs that were highly abundant in extraction and PCR blanks, indicating reagent-derived contamination (Fig. S5). These contaminants included the most abundant OTU in the dataset, *Cladosporium*, a fungus previously associated with insects, including mosquitoes (Accoti et al., 2021; Flores et al., 2022; Hegde et al., 2024) (Fig. S6). After decontamination, we retained 508 OTUs, classified into 77 orders and 320 genera (Supplementary File 3: Table d). We used the ITS2/COI read number ratio as a semi-quantitative measure of fungal load (Fig. 5a). Ratios were generally low (average = 0.04, rarely exceeding 0.5), suggesting limited fungal DNA in mosquito samples. Community composition was highly variable across individuals, locations, and sampling years. The most abundant taxon, *Mrakia* (Basidiomycota: Cystofilobasidiales), accounted for only 8.2% of total fungal reads and was present in 14% of *Ochlerotatus* libraries. The most frequently detected genera were *Monilinia* (Ascomycota:Helotiales) (prevalence 21.7%), and *Phaeosphaeria* (Ascomycota:Pleosporales) (prevalence 20.6%) (Fig. 5b).

**Fig. 5.**
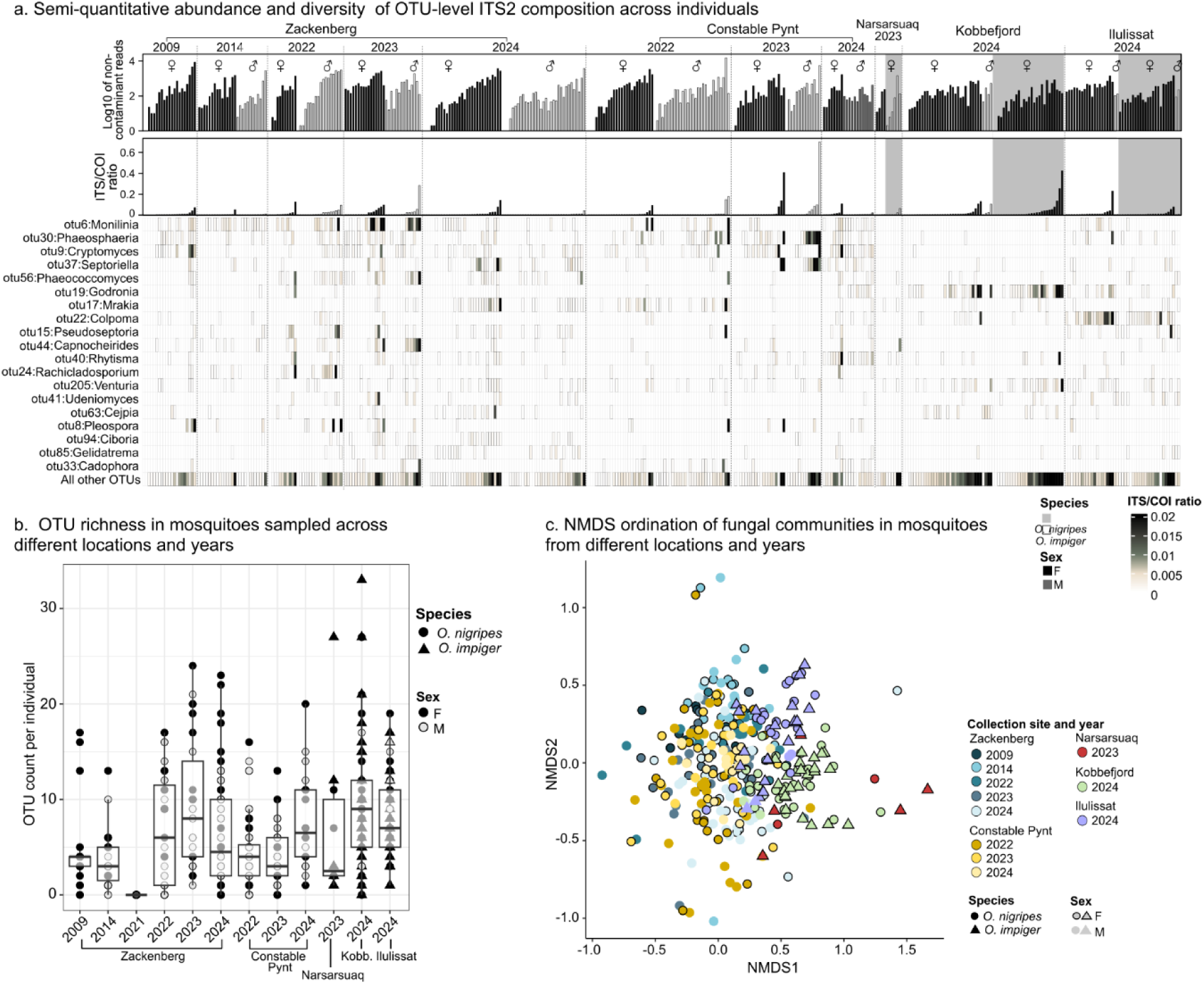
Fungal diversity and distribution across *Ochlerotatus* individuals and populations. a) Fungal OTU richness across sampled populations. Each point represents the number of OTUs in an individual mosquito, with color indicating its sex and shape indicating its species. b) The distribution of fungal abundance and most common OTUs in individual mosquitoes. Barplots indicate the decontaminated total number of ITS2 copies across individuals from the sampled populations (upper panel) and the ITS2/COI ratio per library (bottom panel). The heatmap indicates the abundance of selected fungal OTUs in individual mosquitoes, expressed as ITS2/COI read number ratio. c) NMDS based on Bray-Curtis distances showing females and males of both *Ochlerotatus* species across sampled populations.

Individual libraries retained a median of 234 reads (average 1,240; range 0-29,963), corresponding to an average of 7 OTUs (median 6, range 0-33) (Fig. 5b). NMDS ordination did not fully converge despite 1,000 iterations, which we interpret as further evidence of the high variability and lack of consistent community structure among mosquito individuals. Nonetheless, the ordination revealed moderate differences in fungal communities across locations and years (Fig. 5c), with minimal temporal variation at Constable Pynt. In terms of spatial differences, there was a high degree of overlap in microbial community composition between sites, but fungal communities from East Greenland differed markedly from those in the South/West populations. Among the potential taxa contributing to these differences were diverse plant pathogens, including *Monilinia, Phaeosphaeria, Cryptomyces* (Ascomycota:Rhytismatales)*, and Septoriella*, which were frequent in Zackenberg and Constable Pynt populations*. Godronia* (Ascomycota:Helotiales) and *Colpoma* (Ascomycota:Rhytismatales) were more frequent in Kobbefjord and Ilulissat, respectively. Thus, while some geographic trends were detected, communities within sites remained inconsistent over time (Fig. S7).

### 3.5. Drivers of variation in microbial communities

#### 3.5.1. Bacterial communities

The HMSC models showed a consistent pattern for bacteria: the presence-absence model had moderate explanatory (mean AUC = 0.778) and predictive power under cross-validation (mean AUC = 0.616), whereas the abundance model conditional on presence had high explanatory power (mean R² = 0.80) but no predictive power (mean R² = 0.00). Overall, our predictors explained 16.5% and 93.5% of the variance in the presence-absence and abundance models.

Host individual identity accounted for the largest proportion of explained variation in both models, contributing 8.7% of the total variation to the occurrence model and 72.3% in the abundance model. On the other hand, spatio-temporal factors contributed relatively little, accounting for only 1.9%-2.3% of variation in occurrence and 4.4-5% in abundance variation, while host-specific traits (sex, haplotype, and species) each explained <5% of the variance each in either model (Fig. 6a-b, S8; Tables S4–S5).

**Fig. 6.**
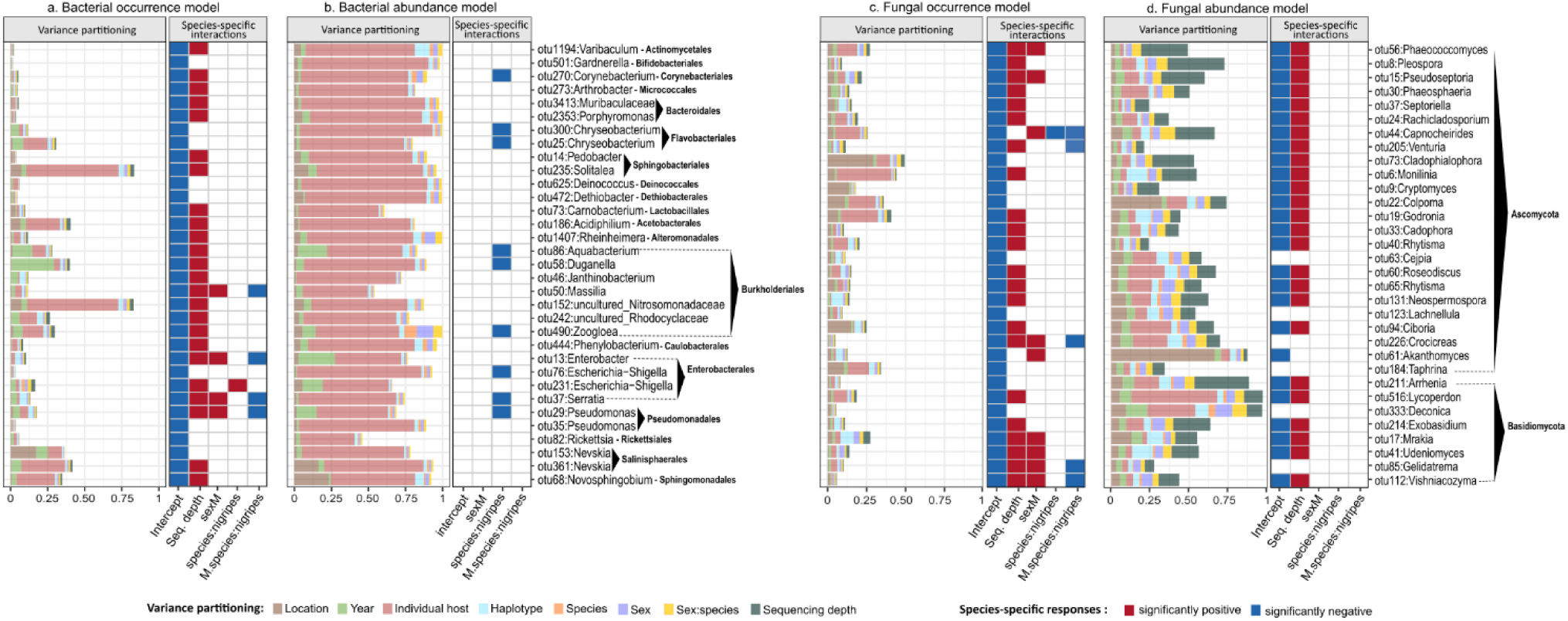
Variance partitioning and responses of the most common bacterial and fungal OTUs to biological covariates. The left panels (a-b) show bacterial OTUs occurring in at least 10% of samples, while the right panels (c-d) show fungal OTUs occurring in at least 5% of samples. Panels a) and c) show results for the presence/absence model, panels b) and d) for the model of abundance conditional on presence. Variance partitioning: the colors represent the proportion of variation attributed to each variable accounted for in the model. Species-specific responses (Beta coefficients): the columns show the posterior probability that the effect in question deviates from zero, with red tiles indicating positive effects, blue tiles indicating negative effects, and white tiles indicating responses that did not gain posterior support with at least 0.95 probability to their direction. Note that given how the model is parameterized, parameter estimates relate to the difference between the factor in question and the reference group (intercept, i.e., females of *O. impiger*). Thus, the colours refer to differences between this group and males of *O. impiger* (sexM); females of *O. nigripes* (species:nigripes), and the interaction sex×species (sexM.species:nigripes).

To assess the systematic drivers of microbial presence/absence, we examined the species-specific responses of microbes to sequencing depth, and to the sex and the species of the individual (Fig. 6a; Fig S9a, Table S6). Most taxa (63.3%) were significantly more likely to be present (or detected) the higher the sequencing depth (Fig. 6a-b; Fig. S6a). A low proportion of taxa (2.2%) occurred with a significantly higher probability in males than in females (Fig. 6a-b; Fig. S9a), and a small minority of common taxa (0.9%, some of them listed in Fig. 6a-b and Fig. S9; i.e., *Aurelimonas*, *Escherichia-Shigella* and *Spiroplasma*) differed in occurrence among mosquito species. Patterns in the abundance model were similarly weak; only_14.6% of OTUs displayed significant species-specific responses, being more abundant in *O. impiger* than in *O. nigripes*, whereas no other predictors showed consistent effects across bacterial taxa (Fig. 6b; Fig. S9b)

To assess potential residual associations among bacterial taxa beyond the effects of measured covariates, we examined OTU–OTU associations from the HMSC models after accounting for fixed and random effects. Residual associations were evaluated at the levels of host individual, haplotype, location, and year (Fig. S10). Overall, statistically significant associations were relatively rare, but among the most abundant and prevalent OTUs, several clear patterns emerged. In the occurrence model, associations varied by hierarchical level, with mostly positive associations at the host-individual and year levels, more negative associations at the location level, and no supported associations at the haplotype level. In the abundance conditional on presence model, supported associations were generally rare and predominantly positive, with the strongest signal at the host-individual level, where high abundance of one taxon tended to coincide with high abundance of others (Fig. S10).

#### 3.5.2 Fungal communities

The HMSC presence-absence model for fungal taxa achieved high explanatory power (mean AUC = 0.88) and moderate predictive power (mean AUC = 0.75), while the abundance conditional on presence model achieved moderate explanatory power (mean R2 = 0.69) but weak predictive power (mean R2 = 0.09). Overall, the measured predictors explained 17.2% of the variation in fungal occurrence and 62.3% of the variation in fungal abundance.

As observed for bacterial communities, host individual identity accounted for the largest proportion of explained variation in both the occurrence (5.2%) and abundance (15.5%) models (Fig. 6b). In the occurrence model, none of the predictors explained more than 5% of the variation. In contrast, in the abundance model, the largest contributions after host individual identity were made by location (7.2%), haplotype (6.7%), and year (6.0%) (Tables S4–S5, S7; Fig. S11).

Species-specific responses showed a strong effect of sequencing depth, with more than half of fungal taxa (55%) showing higher detection probabilities and greater abundances in samples with higher sequencing depth (Table S8; Fig. S12a). Approximately 18% of OTUs were significantly more likely to occur in male *O. impiger* than in male *O. nigripes* (Fig. 6b; included taxa listed in Fig. 5b, such as *Phaeococcomyces, Pseudoseptoria, Capnocheirides*, etc.). Nevertheless, fungal abundance patterns were largely consistent between mosquito species and sexes, with most OTUs showing no significant specific responses (Fig. 6b, Table S8).

Like the bacterial models, the HMSC presence-absence model for fungal taxa achieved moderate predictive power (mean AUC = 0.75) and high explanatory power (mean AUC = 0.88), while the abundance conditional on presence model achieved moderate explanatory power (mean R2 = 0.69) and weak predictive power (mean R2 = 0.09). Overall, only 17.2% of variance was explained by our predictors in the occurrence model and 62.3% in the abundance model.

As observed for bacterial communities, host individual accounted for the largest proportion of explained variation in both occurrence (5.2%) and abundance (15.5%) models (Fig. 6c-d, Table S4). In the occurrence model none of the predictors explained more than 10% of variance (Tables S4–S5; Fig. S11). In contrast, in the abundance conditional on presence model, the largest contributions after host individual identity were by the sampling location (7.2%), COI haplotype (6.7%), and year (6%) (Tables S4–S5; Fig. S11). Over half of the fungal taxa (55%) were more likely to be detected and more abundant as the sequencing depth increased (Fig. 6c-d; Fig. S12a). Approximately 18% of OTUs were significantly more likely to occur in male *O. impiger* than in male *O. nigripes* (Fig. 6d; including taxa such as *Phaeococcomyces, Pseudoseptoria, Capnocheirides*, etc.). Nevertheless, fungal abundance patterns were largely consistent between mosquito species and sexes, with most OTUs showing no significant specific responses (Fig. 6d, Table S8).

After accounting for fixed and random effects in the HMSC models, across host individual, haplotype, location, and year levels (Fig. S13), statistically supported associations were relatively rare. In the occurrence model, positive associations were most frequent at the host-individual level, whereas associations were mixed across locations and absent at the year and haplotype levels. In the abundance conditional on presence model, supported associations were scarce across all levels (Fig. S13).

## 4. Discussion

In this study, we reconstructed the host mitochondrial diversity and geographic distribution of Greenland *Ochlerotatus* mosquitoes sampled across a broad geographic range. In addition, we reconstructed the abundance and composition of both bacterial and fungal microbiota, searching for potential drivers of variation in mosquito-microbe associations. We show that microbial associations in Greenland *Ochlerotatus* mosquitoes are generally sparse and dynamic, with little evidence for widespread, consistent host-associated microbial associations.

### 4.1. Several COI haplotypes mapping two mosquito species in Greenland

Global Biodiversity Information Facility (GBIF) records indicate that *O. impiger* and *O. nigripes* occur across Northern North America, Greenland, and Scandinavia. However, given their phylogenetic proximity and morphological similarity, it is difficult to differentiate between the two species based on morphology alone (Dahl, 2015; Villeneuve et al., 2024). In contrast, our 418 bp COI barcode sequences enabled clear differentiation between the two species, providing valuable insights into their distribution in Greenland.

Within each species, a dominant haplotype was accompanied by several low-frequency variants, often differing by only a single nucleotide. This pattern resulted in low nucleotide diversity despite the presence of multiple haplotypes and is consistent with observations from other mosquito species, including *Aedes albopictus,* where one or a few dominant haplotypes are typically accompanied by numerous rare variants (Francuski et al., 2020; Ismail et al., 2017; Petersen et al., 2024). Such patterns have generally been interpreted as evidence of recent demographic expansion from a limited set of maternal lineages, as in *A. albopictus* (Ismail et al., 2017). In Greenland, the low mitochondrial nucleotide variation combined with species-specific distribution patterns could reflect postglacial colonisation, founder effects, and subsequent local expansion (Porretta et al., 2012).

Beyond the dominant haplotype (median 98% of reads), a few mosquitoes carried low-abundance variants representing >10% of the total COI reads, while most typically carry them in a smaller proportion (< 1% per library). We consider the former as potential cases of heteroplasmy, and the majority of the latter as likely numts (Mastrantonio et al., 2016; Paduan & Ribolla, 2008; Toews & Brelsford, 2012). Interestingly, some of these variants matched the other *Ochlerotatus* species, hinting at incomplete lineage sorting or occasional hybridization, though these interpretations remain tentative without genome-level data.

The non-mosquito COI signal provides an interesting additional window into the biological interactions of these insects with putative parasites and sources of bloodmeal. While the human COI signal in mosquitoes needs to be interpreted carefully due to potential laboratory contamination (and hence we did not consider libraries where the human signal comprised ≤ 12 reads), the presence of other Greenland mammal COI reads is indicative of a feeding interaction. Diet reconstruction using metabarcoding has proven a powerful approach to reconstructing the ecology of wild animals including mosquitoes (Casey et al., 2019; García-Robledo et al., 2013; Vieira et al., 2024), and our data suggest that this information can be obtained alongside wild mosquito identification and microbiome characterisation.

### 4.2. A limited role of bacteria in adult mosquito biology?

While microbes are important or even essential for many blood-feeding insects (Bing et al., 2017; Husnik, 2018; Zhong et al., 2007), our findings suggest a limited role for bacteria in adult Greenland *Ochlerotatus* biology. Compared with data from the present and previous studies on *Aedes aegypti*, *Culex pipiens*, and *Anopheles gambiae* (Habtewold et al., 2016; Villegas et al., 2023), bacterial loads in *Ochlerotatus* species were remarkably low. Our estimated median of ∼8,000 bacterial 16S rRNA copies per mosquito may correspond to only ∼1,000 bacterial cells, depending on rRNA operon copy number, cell ploidy, and genome replication state (Klappenbach, 2001; Louca et al., 2018; Pecoraro et al., 2011). These factors vary across taxa and physiological states, and we may still overestimate true bacterial abundances.

Although cumulative OTU richness across all samples was high, most individual mosquitoes carried only a small subset of these taxa, indicating that much of the observed diversity reflects turnover among hosts rather than consistently maintained associations. Similar patterns have been reported in other mosquitoes, where relatively few taxa are often detected per individual despite substantial among-host variation. For example, field-collected *Anopheles gambiae* showed pronounced inter-individual variation in OTU richness (13-340 OTUs), laboratory-reared *A. aegypti* typically harbored fewer than 20 OTUs, and across multiple mosquito species only 5-10 dominant OTUs are often recovered per individual (Boissière et al., 2012; Muturi et al., 2018; Villegas et al., 2023). Thus, low effective richness per individual is not unique to Greenland *Ochlerotatus* (Fig. S14), but in our system, we demonstrated that it coincides with low bacterial loads and little evidence of stable, recurring bacterial associations.

This interpretation is also supported by the identity of the most common bacteria. *Janthinobacterium, Pseudomonas, Serratia, Enterobacter, Escherichia,* and *Carnobacterium* are all widespread genera that occur across diverse environments and hosts, including other mosquito species (Table S7). For example, *Serratia*, *Carnobacterium*, and *Pseudomonas* were all common and broadly distributed among Swedish phorid flies (Nowak et al., 2025), suggesting that they may represent environmentally widespread, transient bacteria rather than mosquito-specific symbionts. Although some of these bacterial clades can influence nutrition, pathogen susceptibility, or host fitness in other mosquito species, or developmental stages (Guégan et al., 2018), in Greenland *Ochlerotatus*, their low abundance and inconsistent occurrence, even within populations, suggest a limited role in most adult hosts. On the other hand, high bacterial abundance in some individuals suggests cases where microbial effects are more likely to be substantial.

### 4.3. An even smaller role for fungi?

Paralleling the patterns observed in bacteria, fungal communities in Greenland mosquitoes also showed large variation among individuals. Fungal abundances were markedly low overall, with ITS/COI read number ratios generally well below 0.5. While the unknown relative amplification efficiency of these alternative targets precludes absolute abundance estimates, our findings align with data for *A. aegypti*, *C. quinquefasciatus,* and *A. albopictus*, where fungi account for less than half of sequencing reads when both host and fungal signal are co-amplified (Hegde et al., 2024).

The low numbers of fungal OTUs we observed in Ochlerotatus mosquitoes (median 7, range 0-30) are also comparable to those observed in other species. For example, wild *C. quinquefasciatus* have been reported to harbor ∼10-20 fungal OTUs, *A. aegypti* and *A. albopictus* ∼10-15 (Hegde et al., 2024; Thongsripong et al., 2018), while in laboratory-reared *C. quinquefasciatus,* the number of OTUs ranged between 5 and 22 (Flores et al., 2022).

Community composition was dominated by a handful of taxa detected across sites, with higher prevalence at specific locations, and no OTU was consistently shared among populations. Thus, as with bacteria, we found no evidence for a core fungal microbiome. Such variation in microbiota among hosts matches patterns previously described in *C. pipiens* and *Culiseta incidens*, where only a few microbial genera recur, without clear patterns in prevalence (Chandler et al., 2015).

Similarly to bacteria, most fungal OTUs detected in *Ochlerotatus* also appear to be environmentally derived. The most prevalent taxa include genera commonly associated with plants and other terrestrial substrates, such as *Cryptomyces* and *Septoriella*, while several others (e.g. *Mrakia*, *Phaeosphaeria*, *Godronia*) have also been detected in Greenland environmental DNA surveys (Crous et al., 2015; Lantz et al., 2011; Ovaskainen et al., 2020). Although some of these genera have occasionally been reported from animals (Alexiev et al., 2021; De Miccolis Angelini et al., 2022; McArt et al., 2016), their taxonomic identities and known ecologies suggest that they are more likely acquired from the surrounding environment than maintained as persistent mosquito associates. Overall, fungi appear to play an even less consistent role than bacteria in adult Greenland mosquitoes. Importantly, our study was focused on adult mosquitoes alone. Thus, our current findings suggest that bacterial and fungal communities are unlikely to play major roles in adult Greenland mosquitoes given their inconsistency across locations and time. For mosquito larvae, microbial diversity might well have a different importance (Ravenscraft & Coon, 2026), leaving ample scope for future work.

### 4.4. The mosquito microbiome: patterns, drivers, and likely consequences

The HMSC results further support the view that microbial associations in Greenland *Ochlerotatus* are weakly structured and largely context dependent. Host-related factors (haplotype, species, sex) and spatio-temporal drivers explained little variation in either overall bacterial or fungal community composition or in the occurrence of individual OTUs, suggesting that much of the observed variation is driven either by stochastic processes or by factors not accounted for in our models. Because mosquitoes develop in aquatic habitats and subsequently interact with a variety of environmental surfaces and food sources as adults, it is likely that many of the detected microorganisms represent taxa from larval habitats and acquisitions from nectar sources, resting sites, or vertebrate hosts (Moradi-Asl & Soltan-Alinejad, 2026). Although the abundance model for both bacteria and fungi showed high explanatory power, their poor predictive performance under cross-validation suggests that these abundance patterns were not robustly generalizable. In bacterial communities, variation among individual mosquitoes represented the dominant source of explained variation, possibly reflecting differences in physiology, immune conditions, or individual life histories. In fungal communities, the absence of spike-in correction combined with low fungal biomass resulted in sequencing depth explaining abundance variation in a similar manner as individual host. Nevertheless, sampling location had an effect on the abundance of some taxa, including *Akanthomyces*, a genus encompassing known entomopathogens of invertebrates (Aini et al., 2020), and *Colpoma*, a saprophytic fungus associated with trees that has occasionally been isolated from invertebrates (Gierczyk et al., 2019; S. Lee et al., 2016). Notably, the presence of such taxa adds an additional layer of complexity to the task of disentangling what drives the estructure of microbial communities.

Overall, these microbial diversity patterns attest to a largely environmental origin and stochastic acquisition of microbes in our system. Among the few mosquito-associated bacteria showing responses to the spatio-temporal drivers, *Moheibacter*, *Duganella*, *Neochlamydia*, *Nevskia*, *Shewanella*, and members of the order Acetobacterales are typically associated with soil and aquatic environments (Choi et al., 2015; Liu et al., 2021; Roman et al., 2025). Similarly, the fungi *Cryptomyces*, *Ciboria*, *Cladophialophora*, *Juncaceicola*, and *Everhartia* are primarily environmental taxa, commonly associated with plant material, soil, or other terrestrial substrates (Badali et al., 2008; Becarelli et al., 2021; Lantz et al., 2011; Ovaskainen et al., 2020). Patterns of residual covariation among microbial taxa (Fig S10, S13) further support this interpretation. Some bacterial and fungal taxa tended to co-occur, or to reach high abundance in the same mosquitoes, without corresponding shifts across the rest of the microbial community. Rather than indicating tightly integrated microbial consortia in adult *Ochlerotatus*, such associations may instead reflect shared environmental sources, similar exposure histories, or parallel responses to within-host ecological opportunities. In this sense, individual mosquitoes may be sampling different subsets of locally available microbes, with only some taxa occasionally persisting or proliferating after acquisition.More broadly, our findings for *O. nigripes* and *O. impiger* support the view that not all animals necessarily maintain a stable or functionally meaningful microbiome, and many carry sparse and inconsistent assemblages dominated by transient environmental taxa, occasional pathogens, or opportunistic colonists (Hammer et al., 2019; Ravenscraft & Coon, 2026). In the apparent absence of heritable intracellular symbionts or a consistent extracellular core microbiota, *Ochlerotatus* adult mosquitoes may emerge virtually sterile from the pupal stage. Subsequently, they acquire both bacteria and fungi from resting substrates and through nectar or blood feeding. The resulting microbiome would then depend strongly on local exposure, stochastic colonisation, and the ability of individual taxa to persist or grow within the mosquito.

Under this view, the striking variation observed among individuals in microbial abundance and diversity may reflect differences in age, feeding history, resting behavior, immune competence, or within-host microbial competition. Some mosquitoes may simply have had more opportunities for environmental acquisition. Whether such variation is largely incidental or whether some environmentally acquired bacteria or fungi can still contribute meaningfully to host physiology under specific ecological contexts remains an important question for future work.

## 5. Conclusions

While blood-feeding insects will often rely on specialised microbiota to survive on their unbalanced diet, our study offers no evidence for such a core community in *Ochlerotatus nigripes* and *O. impiger* from across Greenland. Rather, it highlights extensive variation at the individual level. Greenland mosquitoes appear to form opportunistic and typically low-abundance associations with diverse bacteria and fungi representing clades broadly distributed in the environment. From the perspective of mutualistic interactions, Arctic mosquitoes can thus be described as ’cold and lonely’, a fitting description of insects that are not only adapted to thrive in a frigid, inhospitable environment but also lack abundant host-adapted microbes, surviving in isolation on a challenging diet. Needless to say, we cannot rule out the possibility that some of the identified microbes will still significantly affect insect performance, even if it seems unlikely that they are critical to successful growth and reproduction. With the growing awareness of the diversity of microbial associations and the absence of specialised microbiota in many insect clades, and the improving methodological and conceptual toolkit for microbiota study, we are in a good position to assess how common such patterns may be among other mosquitoes and insects and their relevance for the host biology.

## Acknowledgements

We thank Ewa Chrostek and the members of the Chrostek lab for providing the *Ae. aegypti* specimens and for generously offering their facilities, resources, and expertise for the rearing and maintenance of the specimens.This project was supported by the Polish National Agency for Academic Exchange grant no. PPN/PPO/2018/1/00015, and Polish National Science Centre grants no. 2018/31/B/NZ8/01158 and 2021/43/B/NZ8/03376 and has received funding from INTERACT III Transnational Access under the European Union H2020 Grant Agreement No. 629 and 729. Sequencing was supported by a grant from the Priority Research Area BioS under the Strategic Programme Excellence Initiative at Jagiellonian University in Kraków and by the National Genomics Infrastructure in Stockholm. The contribution of TR was funded by the European Union: the European Research Council (ERC) under the European Union’s Horizon 2020 research and innovation programme (grant agreement No 856506: ERC-synergy project LIFEPLAN) and the Swedish Research Council *Vetenskapsrådet* (grant 2023-05118).

## Availability of data and materials

The dataset supporting the conclusions of this article is available in the National Centre for Biotechnology Information repository under the BioProject PRJNA1330896 (https://www.ncbi.nlm.nih.gov/bioproject/?term=PRJNA1330896). Raw sequencing read data (i.e., COI, 16S rRNA, and ITS2) obtained from this dataset are available at http://doi.org/10.6084/m9.figshare.c.8012695 and http://doi.org/10.6084/m9.figshare.31934055.

## References

Abarenkov, K., Nilsson, R. H., Larsson, K.-H., Taylor, A. F. S., May, T. W., Frøslev, T. G., Pawlowska, J., Lindahl, B., Põldmaa, K., Truong, C., Vu, D., Hosoya, T., Niskanen, T., Piirmann, T., Ivanov, F., Zirk, A., Peterson, M., Cheeke, T. E., Ishigami, Y., … Kõljalg, U. (2024). The UNITE database for molecular identification and taxonomic communication of fungi and other eukaryotes: Sequences, taxa and classifications reconsidered. Nucleic Acids Research, 52(D1), D791–D797. 10.1093/nar/gkad1039

Aini, A. N., Mongkolsamrit, S., Wijanarka, W., Thanakitpipattana, D., Luangsa-ard, J. J., & Budiharjo, A. (2020). Diversity of Akanthomyces on moths (Lepidoptera) in Thailand. MycoKeys, 71, 1–22. 10.3897/mycokeys.71.55126

Alexiev, A., Chen, M. Y., & McKenzie, V. J. (2021). Identifying fungal-host associations in an amphibian host system. PLOS ONE, 16(8), e0256328. 10.1371/journal.pone.0256328

Alomar, A. A., Pérez-Ramos, D. W., Kim, D., Kendziorski, N. L., Eastmond, B. H., Alto, B. W., & Caragata, E. P. (2023). Native Wolbachia infection and larval competition stress shape fitness and West Nile virus infection in Culex quinquefasciatus mosquitoes. Frontiers in Microbiology, 14, 1138476. 10.3389/fmicb.2023.1138476

Antich, A., Palacin, C., Wangensteen, O. S., & Turon, X. (2021). To denoise or to cluster, that is not the question: Optimizing pipelines for COI metabarcoding and metaphylogeography. BMC Bioinformatics, 22(1). 10.1186/s12859-021-04115-6

Badali, H., Gueidan, C., Najafzadeh, M. J., Bonifaz, A., Van Den Ende, A. H. G. G., & De Hoog, G. S. (2008). Biodiversity of the genus Cladophialophora. Studies in Mycology, 61, 175–191. 10.3114/sim.2008.61.18

Bai, L., Wang, L., Vega-Rodríguez, J., Wang, G., & Wang, S. (2019). A gut symbiotic bacterium Serratia marcescens renders mosquito resistance to Plasmodium Infection through activation of mosquito Immune responses. Frontiers in Microbiology, 10, 1580. 10.3389/fmicb.2019.01580

Bandelt, H. J., Forster, P., & Rohl, A. (1999). Median-joining networks for inferring intraspecific phylogenies. Molecular Biology and Evolution, 16(1), 37–48. 10.1093/oxfordjournals.molbev.a026036

Barredo, E., & DeGennaro, M. (2020). Not Just from Blood: Mosquito Nutrient Acquisition from Nectar Sources. Trends in Parasitology, 36(5), 473–484. 10.1016/j.pt.2020.02.003

Becarelli, S., Chicca, I., La China, S., Siracusa, G., Bardi, A., Gullo, M., Petroni, G., Levin, D. B., & Di Gregorio, S. (2021). A new Ciboria sp. For soil mycoremediation and the bacterial contribution to the depletion of total petroleum hydrocarbons. Frontiers in Microbiology, 12, 647373. 10.3389/fmicb.2021.647373

Bhattacharyya, S., Engl, T., Kaltenpoth, M., & Ferrarini, M. G. (2026). Regulation of beneficial intracellular symbionts in insects. Current Opinion in Insect Science, 77, 101566. 10.1016/j.cois.2026.101566

Bing, X., Attardo, G. M., Vigneron, A., Aksoy, E., Scolari, F., Malacrida, A., Weiss, B. L., & Aksoy, S. (2017). Unravelling the relationship between the tsetse fly and its obligate symbiont *Wigglesworthia*: Transcriptomic and metabolomic landscapes reveal highly integrated physiological networks. Proceedings of the Royal Society B: Biological Sciences, 284(1857), 20170360. 10.1098/rspb.2017.0360

Böcher, J. (2015). Chapter 4: The Greenland entomofauna: Zoogeography and history. In The Greenland Entomofauna: An identification manual of insects, spiders and their allies (Vol. 44). Brill.

Boissière, A., Tchioffo, M. T., Bachar, D., Abate, L., Marie, A., Nsango, S. E., Shahbazkia, H. R., Awono-Ambene, P. H., Levashina, E. A., Christen, R., & Morlais, I. (2012). Midgut microbiota of the malaria mosquito vector Anopheles gambiae and interactions with Plasmodium falciparum infection. PLoS Pathogens, 8(5), e1002742. 10.1371/journal.ppat.1002742

Borman, T., Ernst, G., Sudarshan, A. S., & Lahti, L. (2025). mia: Microbiome analysis [Github]. https://github.com/microbiome/mia

Buczek, M., Kolasa, M. R., Prus-Frankowska, M., Lipowska, M., Nowak, K. H., Azarbad, H., Franco, D. C., Marszałek, M., Roslin, T., Michalik, A., & Łukasik, P. (2024). A new tool in a toolbox: Addressing challenges in high-throughput microbiota surveys across diverse wild insects. 10.1101/2024.08.26.609764

Casey, J. M., Meyer, C. P., Morat, F., Brandl, S. J., Planes, S., & Parravicini, V. (2019). Reconstructing hyperdiverse food webs: Gut content metabarcoding as a tool to disentangle trophic interactions on coral reefs. Methods in Ecology and Evolution, 10(8), 1157–1170. 10.1111/2041-210X.13206

Chandler, J. A., Liu, R. M., & Bennett, S. N. (2015). RNA shotgun metagenomic sequencing of northern California (USA) mosquitoes uncovers viruses, bacteria, and fungi. Frontiers in Microbiology, 06. 10.3389/fmicb.2015.00185

Choi, S. Y., Kim, S., Lyuck, S., Kim, S. B., & Mitchell, R. J. (2015). High-level production of violacein by the newly isolated Duganella violaceinigra str. NI28 and its impact on Staphylococcus aureus. Scientific Reports, 5(1), 15598. 10.1038/srep15598

Clement, M., Snell, Q., Walke, P., Posada, D., & Crandall, K. (2002). TCS: Estimating gene genealogies. Proceedings 16th International Parallel and Distributed Processing Symposium, 7 pp. 10.1109/IPDPS.2002.1016585

Coon, K. L., Brown, M. R., & Strand, M. R. (2016). Mosquitoes host communities of bacteria that are essential for development but vary greatly between local habitats. Molecular Ecology, 25(22), 5806–5826. 10.1111/mec.13877

Coon, K. L., Vogel, K. J., Brown, M. R., & Strand, M. R. (2014). Mosquitoes rely on their gut microbiota for development. Molecular Ecology, 23(11), 2727–2739. 10.1111/mec.12771

Corbet, P. S. (1967). Facultative autogeny in Arctic mosquitoes. Nature, 215(5101), 662–663. 10.1038/215662a0

Corbet, P. S., & Downe, A. E. R. (1966). Natural hosts of mosquitoes in Northern Ellesmere island. ARCTIC, 19(2), 153–161. 10.14430/arctic3422

Crous, P. W., Carris, L. M., Giraldo, A., Groenewald, J. Z., Hawksworth, D. L., Hemández-Restrepo, M., Jaklitsch, W. M., Lebrun, M.-H., Schumacher, R. K., Stielow, J. B., Van Der Linde, E. J., Vilcāne, J., Voglmayr, H., & Wood, A. R. (2015). The genera of fungi - fixing the application of the type species of generic names - G 2: Allantophomopsis, Latorua, Macrodiplodiopsis, Macrohilum, Milospium, Protostegia, Pyricularia, Robillarda, Rotula, Septoriella, Torula, and Wojnowicia. IMA Fungus, 6(1), 163–198. 10.5598/imafungus.2015.06.01.11

Dahl, C. (2015). Chapter 17. Diptera (Two-winged or “true” flies). In The Greenland Entomofauna: An identification manual of insects, spiders and their allies (Vol. 44). Brill.

De Miccolis Angelini, R. M., Landi, L., Raguseo, C., Pollastro, S., Faretra, F., & Romanazzi, G. (2022). Tracking of diversity and evolution in the brown rot fungi Monilinia fructicola, Monilinia fructigena, and Monilinia laxa. Frontiers in Microbiology, 13, 854852. 10.3389/fmicb.2022.854852

Douglas, A. E. (2022). Insects and their beneficial microbes. Princeton University Press. 10.2307/j.ctv24q54q5

Edgar, R. C. (2010). Search and clustering orders of magnitude faster than BLAST. Bioinformatics, 26(19), 2460–2461. 10.1093/bioinformatics/btq461

Elbrecht, V., Braukmann, T. W. A., Ivanova, N. V., Prosser, S. W. J., Hajibabaei, M., Wright, M., Zakharov, E. V., Hebert, P. D. N., & Steinke, D. (2019). Validation of COI metabarcoding primers for terrestrial arthropods. PeerJ, 7, e7745. 10.7717/peerj.7745

Flores, G. A. M., Lopez, R. P., Cerrudo, C. S., Consolo, V. F., & Berón, C. M. (2022). Culex quinquefasciatus holobiont: A fungal metagenomic approach. Frontiers in Fungal Biology, 3, 918052. 10.3389/ffunb.2022.918052

Francuski, L., Ludoški, J., Milutinović, A., Krtinić, B., & Milankov, V. (2020). Comparative phylogeography and integrative taxonomy of Ochlerotatus caspius (Dipera: Culicidae) and Ochlerotatus dorsalis. *Journal of Medical Entomology*, tjaa153. 10.1093/jme/tjaa153

García-Robledo, C., Erickson, D. L., Staines, C. L., Erwin, T. L., & Kress, W. J. (2013). Tropical plant–herbivore networks: Reconstructing species interactions using DNA barcodes. PLoS ONE, 8(1), e52967. 10.1371/journal.pone.0052967

Gierczyk, B., Szczepkowski, A., Ślusarczyk, T., & Kujawa, A. (2019). Contribution to knowledge of fungal biota of Kampinos National Park (Poland): Part 3. Acta Mycologica, 54(2). 10.5586/am.1129

Gu, Z. (2022). Complex heatmap visualization. iMeta, 1(3), e43. 10.1002/imt2.43

Gu, Z., Eils, R., & Schlesner, M. (2016). Complex heatmaps reveal patterns and correlations in multidimensional genomic data. Bioinformatics, 32(18), 2847–2849. 10.1093/bioinformatics/btw313

Guégan, M., Zouache, K., Démichel, C., Minard, G., Tran Van, V., Potier, P., Mavingui, P., & Valiente Moro, C. (2018). The mosquito holobiont: Fresh insight into mosquito-microbiota interactions. Microbiome, 6(1), 49. 10.1186/s40168-018-0435-2

Habtewold, T., Duchateau, L., & Christophides, G. K. (2016). Flow cytometry analysis of the microbiota associated with the midguts of vector mosquitoes. Parasites & Vectors, 9(1), 167. 10.1186/s13071-016-1438-0

Hall, M. (2022). Mosquitopia: The place of pests in a healthy world (D. Tamir, Ed.). Taylor & Francis.

Hammer, T. J., Janzen, D. H., Hallwachs, W., Jaffe, S. P., & Fierer, N. (2017). Caterpillars lack a resident gut microbiome. Proceedings of the National Academy of Sciences, 114(36), 9641–9646. 10.1073/pnas.1707186114

Hammer, T. J., Sanders, J. G., & Fierer, N. (2019). Not all animals need a microbiome. FEMS Microbiology Letters, 366(10), fnz117. 10.1093/femsle/fnz117

Hegde, S., Khanipov, K., Albayrak, L., Golovko, G., Pimenova, M., Saldaña, M. A., Rojas, M. M., Hornett, E. A., Motl, G. C., Fredregill, C. L., Dennett, J. A., Debboun, M., Fofanov, Y., & Hughes, G. L. (2018). Microbiome interaction networks and community structure from laboratory-reared and field-collected Aedes aegypti, Aedes albopictus, and Culex quinquefasciatus mosquito vectors. Frontiers in Microbiology, 9, 2160. 10.3389/fmicb.2018.02160

Hegde, S., Khanipov, K., Hornett, E. A., Nilyanimit, P., Pimenova, M., Saldaña, M. A., De Bekker, C., Golovko, G., & Hughes, G. L. (2024). Interkingdom interactions shape the fungal microbiome of mosquitoes. Animal Microbiome, 6(1), 11. 10.1186/s42523-024-00298-4

Hegde, S., Rasgon, J. L., & Hughes, G. L. (2015). The microbiome modulates arbovirus transmission in mosquitoes. Current Opinion in Virology, 15, 97–102. 10.1016/j.coviro.2015.08.011

Hoffmann, A. A., Ahmad, N. W., Keong, W. M., Ling, C. Y., Ahmad, N. A., Golding, N., Tierney, N., Jelip, J., Putit, P. W., Mokhtar, N., Sandhu, S. S., Ming, L. S., Khairuddin, K., Denim, K., Rosli, N. M., Shahar, H., Omar, T., Ridhuan Ghazali, M. K., Aqmar Mohd Zabari, N. Z., … Sinkins, S. P. (2024). Introduction of Aedes aegypti mosquitoes carrying wAlbB Wolbachia sharply decreases dengue incidence in disease hotspots. iScience, 27(2), 108942. 10.1016/j.isci.2024.108942

Husnik, F. (2018). Host–symbiont–pathogen interactions in blood-feeding parasites: Nutrition, immune cross-talk and gene exchange. Parasitology, 145(10), 1294–1303. 10.1017/s0031182018000574

Ismail, N.-A., Adilah-Amrannudin, N., Hamsidi, M., Ismail, R., Dom, N. C., Ahmad, A. H., Mastuki, M. F., & Camalxaman, S. N. (2017). The genetic diversity, haplotype analysis, and phylogenetic relationship of Aedes albopictus (Diptera: Culicidae) based on the cytochrome oxidase 1 marker: a Malaysian scenario. Journal of Medical Entomology, 54(6), 1573–1581. 10.1093/jme/tjx126

Iwaszkiewicz-Eggebrecht, E., Łukasik, P., Buczek, M., Deng, J., Hartop, E. A., Havnås, H., Prus-Frankowska, M., Ugarph, C. R., Viteri, P., Andersson, A. F., Roslin, T., Tack, A. J. M., Ronquist, F., & Miraldo, A. (2023). FAVIS: Fast and versatile protocol for non-destructive metabarcoding of bulk insect samples. PLOS ONE, 18(7), e0286272. 10.1371/journal.pone.0286272

Klappenbach, J. A. (2001). rrndb: The ribosomal RNA operon copy number database. Nucleic Acids Research, 29(1), 181–184. 10.1093/nar/29.1.181

Lahti, L., & Sudarshan, A. S. (2017). Tools for microbiome analysis in R. http://microbiome.github.com/microbiome

Lange, C., Boyer, S., Bezemer, T. M., Lefort, M.-C., Dhami, M. K., Biggs, E., Groenteman, R., Fowler, S. V., Paynter, Q., Verdecia Mogena, A. M., & Kaltenpoth, M. (2023). Impact of intraspecific variation in insect microbiomes on host phenotype and evolution. The ISME Journal, 17(11), 1798–1807. 10.1038/s41396-023-01500-2

Lantz, H., Johnston, P. R., Park, D., & Minter, D. W. (2011). Molecular phylogeny reveals a core clade of Rhytismatales. Mycologia, 103(1), 57–74. 10.3852/10-060

Lee, M., O’Rorke, R., Clark, N. J., Webster, T. U., & Wells, K. (2025). Diversity and Plasticity in Mosquito Feeding Patterns: A Meta-Analysis of ‘Universal’ DNA Diet Studies. Global Ecology and Biogeography, 34(6), e70077. 10.1111/geb.70077

Lee, S., Tamayo-Castillo, G., Pang, C., Clardy, J., Cao, S., & Kim, K. H. (2016). Diketopiperazines from Costa Rican endolichenic fungus Colpoma sp. CR1465A. Bioorganic & Medicinal Chemistry Letters, 26(10), 2438–2441. 10.1016/j.bmcl.2016.03.115

Leigh, J. W., & Bryant, D. (2015). POPART: Full-feature software for haplotype network construction. Methods in Ecology and Evolution, 6(9), 1110–1116. 10.1111/2041-210X.12410

Leray, M., Knowlton, N., & Machida, R. J. (2022). MIDORI2: A collection of quality controlled, preformatted, and regularly updated reference databases for taxonomic assignment of eukaryotic mitochondrial sequences. Environmental DNA, 4(4), 894–907. 10.1002/edn3.303

Liu, Y., Chang, Y.-Q., Wang, C.-N., Ye, M.-Q., Wang, M.-Y., & Du, Z.-J. (2021). Moheibacter lacus sp. Nov., Isolated from Freshwater Lake Sediment. Current Microbiology, 78(5), 2160–2164. 10.1007/s00284-021-02465-1

Louca, S., Doebeli, M., & Parfrey, L. W. (2018). Correcting for 16S rRNA gene copy numbers in microbiome surveys remains an unsolved problem. Microbiome, 6(1), 41. 10.1186/s40168-018-0420-9

Łukasik, P., Newton, J. A., Sanders, J. G., Hu, Y., Moreau, C. S., Kronauer, D. J. C., O’Donnell, S., Koga, R., & Russell, J. A. (2017). The structured diversity of specialized gut symbionts of the New World army ants. Molecular Ecology, 26(14), 3808–3825. 10.1111/mec.14140

Malacrinò, A. (2022). Host species identity shapes the diversity and structure of insect microbiota. Molecular Ecology, 31(3), 723–735. 10.1111/mec.16285

Mastrantonio, V., Porretta, D., Urbanelli, S., Crasta, G., & Nascetti, G. (2016). Dynamics of mtDNA introgression during species range expansion: Insights from an experimental longitudinal study. Scientific Reports, 6(1), 30355. 10.1038/srep30355

McArt, S. H., Miles, T. D., Rodriguez-Saona, C., Schilder, A., Adler, L. S., & Grieshop, M. J. (2016). Floral scent mimicry and vector-pathogen associations in a Pseudoflower-Inducing plant pathogen system. PLOS ONE, 11(11), e0165761. 10.1371/journal.pone.0165761

McMurdie, P. J., & Holmes, S. (2013). phyloseq: An R package for reproducible interactive analysis and graphics of microbiome census data. PLoS ONE, 8(4), e61217. 10.1371/journal.pone.0061217

Minwuyelet, A., Petronio, G. P., Yewhalaw, D., Sciarretta, A., Magnifico, I., Nicolosi, D., Di Marco, R., & Atenafu, G. (2023). Symbiotic Wolbachia in mosquitoes and its role in reducing the transmission of mosquito-borne diseases: Updates and prospects. Frontiers in Microbiology, 14, 1267832. 10.3389/fmicb.2023.1267832

Moonen, J. P., Schinkel, M., Van Der Most, T., Miesen, P., & Van Rij, R. P. (2023). Composition and global distribution of the mosquito virome—A comprehensive database of insect-specific viruses. One Health, 16, 100490. 10.1016/j.onehlt.2023.100490

Moradi-Asl, E., & Soltan-Alinejad, P. (2026). Impact of environmental factors on mosquito midgut microbiota. Medical and Veterinary Entomology, mve.70101. 10.1111/mve.70101

Müllerová, J., Elsterová, J., Černý, J., Ditrich, O., Žárský, J., Culler, L. E., Kampen, H., Walther, D., Coulson, S. J., Růžek, D., & Grubhoffer, L. (2018). No indication of arthropod-vectored viruses in mosquitoes (Diptera: Culicidae) collected on Greenland and Svalbard. Polar Biology, 41(8), 1581–1586. 10.1007/s00300-017-2242-9

Muturi, E. J., Lagos-Kutz, D., Dunlap, C., Ramirez, J. L., Rooney, A. P., Hartman, G. L., Fields, C. J., Rendon, G., & Kim, C.-H. (2018). Mosquito microbiota cluster by host sampling location. Parasites & Vectors, 11(1), 468. 10.1186/s13071-018-3036-9

Nowak, K. H., Hartop, E., Prus-Frankowska, M., Buczek, M., Kolasa, M., Roslin, T., Ovaskainen, O., & Łukasik, P. (2025). What lurks in the dark? An innovative framework for studying diverse wild insect microbiota. Microbiome, 13(186). 10.1186/s40168-025-02169-9

Ovaskainen, O. (2020). Joint species distribution modelling: With applications in R. Cambridge University Press.

Ovaskainen, O., Abrego, N., Somervuo, P., Palorinne, I., Hardwick, B., Pitkänen, J.-M., Andrew, N. R., Niklaus, P. A., Schmidt, N. M., Seibold, S., Vogt, J., Zakharov, E. V., Hebert, P. D. N., Roslin, T., & Ivanova, N. V. (2020). Monitoring fungal communities with the global spore sampling project. Frontiers in Ecology and Evolution, 7, 511. 10.3389/fevo.2019.00511

Ovaskainen, O., Tikhonov, G., Norberg, A., Guillaume Blanchet, F., Duan, L., Dunson, D., Roslin, T., & Abrego, N. (2017). How to make more out of community data? A conceptual framework and its implementation as models and software. Ecology Letters, 20(5), 561–576. 10.1111/ele.12757

Paduan, K. D. S., & Ribolla, P. E. M. (2008). Mitochondrial DNA polymorphism and heteroplasmy in populations of Aedes aegypti in Brazil. Journal of Medical Entomology, 45(1), 59–67. 10.1603/0022-2585(2008)45%5B59:MDPAHI%5D2.0.CO;2

Parada, A. E., Needham, D. M., & Fuhrman, J. A. (2016). Every base matters: Assessing small subunit RRNA primers for marine microbiomes with mock communities, time series and global field samples. Environmental Microbiology, 18(5), 1403–1414. 10.1111/1462-2920.13023

Pecoraro, V., Zerulla, K., Lange, C., & Soppa, J. (2011). Quantification of ploidy in Proteobacteria revealed the existence of monoploid, (Mero-)Oligoploid and polyploid species. PLoS ONE, 6(1), e16392. 10.1371/journal.pone.0016392

Petersen, V., Devicari, M., & Suesdek, L. (2015). High morphological and genetic variabilities of Ochlerotatus scapularis, a potential vector of filarias and arboviruses. Parasites & Vectors, 8(1), 128. 10.1186/s13071-015-0740-6

Petersen, V., Santana, M., Karina-Costa, M., Nachbar, J. J., Martin-Martin, I., Adelman, Z. N., & Burini, B. C. (2024). Aedes (Ochlerotatus) scapularis, Aedes japonicus japonicus, and Aedes (Fredwardsius) vittatus (Diptera: Culicidae): three neglected mosquitoes with potential global health risks. Insects, 15(8), 600. 10.3390/insects15080600

Płoszka, Z., Nowak, K. H., Tischer, M., Michalik, A., Kolasa, M. R., & Łukasik, P. (2025). Dissecting multitrophic interactions: The relationships among Entomophthora, their dipteran hosts, and associated bacteria. Journal of Invertebrate Pathology, 213, 108425. 10.1016/j.jip.2025.108425

Porretta, D., Mastrantonio, V., Bellini, R., Somboon, P., & Urbanelli, S. (2012). Glacial history of a modern invader: Phylogeography and species distribution modelling of the Asian Tiger Mosquito Aedes albopictus. PLoS ONE, 7(9), e44515. 10.1371/journal.pone.0044515

Quast, C., Pruesse, E., Yilmaz, P., Gerken, J., Schweer, T., Yarza, P., Peplies, J., & Glöckner, F. O. (2012). The SILVA ribosomal RNA gene database project: Improved data processing and web-based tools. Nucleic Acids Research, 41(D1), D590–D596. 10.1093/nar/gks1219

Ravenscraft, A., & Coon, K. L. (2026). Transient Microbes in Insects: Fleeting but Functional. Annual Review of Entomology, 71(1), 253–273. 10.1146/annurev-ento-121423-013446

Reeves, W. K., Breidenbaugh, M. S., Thomas, E. E., & Glowacki, M. N. (2013). Mosquitoes of Thule air base, Greenland. Journal of the American Mosquito Control Association, 29(4), 383–384. 10.2987/13-6341.1

Rognes, T., Flouri, T., Nichols, B., Quince, C., & Mahé, F. (2016). VSEARCH: A versatile open source tool for metagenomics. PeerJ, 4, e2584. 10.7717/peerj.2584

Rohland, N., & Reich, D. (2012). Cost-effective, high-throughput DNA sequencing libraries for multiplexed target capture. Genome Research, 22(5), 939–946. 10.1101/gr.128124.111

Roman, F. A., Byrne, T., Martin, R. L., Mena-Aguilar, D., Smeltz, R. E., Finkelstein, R., Pruden, A., & Edwards, M. A. (2025). Retrospective analysis of drinking water microcosm microbiomes reveals an apparent antagonistic relationship between Neochlamydia and Legionella. Environmental Science & Technology Letters, 12(8), 990–996. 10.1021/acs.estlett.5c00590

Salter, S. J., Cox, M. J., Turek, E. M., Calus, S. T., Cookson, W. O., Moffatt, M. F., Turner, P., Parkhill, J., Loman, N. J., & Walker, A. W. (2014). Reagent and laboratory contamination can critically impact sequence-based microbiome analyses. BMC Biology, 12(1), 87. 10.1186/s12915-014-0087-z

Sanders, J. G., Łukasik, P., Frederickson, M. E., Russell, J. A., Koga, R., Knight, R., & Pierce, N. E. (2017). Dramatic differences in gut bacterial densities correlate with diet and habitat in rainforest ants. Integrative and Comparative Biology, 57(4), 705–722. 10.1093/icb/icx088

Schilling, M., Jagdev, M., Thomas, H., & Johnson, N. (2025). Metagenomic analysis of mosquitoes from Kangerlussuaq, Greenland reveals a unique virome. Scientific Reports, 15(1), 17141. 10.1038/s41598-025-01086-z

Snyman, J., Villeneuve, C., Snyman, L. P., Martinez, V., Dusfour, I., Lecomte, N., Jenkins, E. J., Hobman, T. C., Leighton, P. A., & Kumar, A. (2024). First confirmed report of Jamestown Canyon Virus in Greenland. Journal of Medical Virology, 96(11), e70064. 10.1002/jmv.70064

Thompson, L. R., Sanders, J. G., McDonald, D., Amir, A., Ladau, J., Locey, K. J., Prill, R. J., Tripathi, A., Gibbons, S. M., Ackermann, G., Navas-Molina, J. A., Janssen, S., Kopylova, E., Vázquez-Baeza, Y., González, A., Morton, J. T., Mirarab, S., Zech Xu, Z., Jiang, L., … Zhao, H. (2017). A communal catalogue reveals Earth’s multiscale microbial diversity. Nature, 551(7681), 457–463. 10.1038/nature24621

Thongsripong, P., Chandler, J. A., Green, A. B., Kittayapong, P., Wilcox, B. A., Kapan, D. D., & Bennett, S. N. (2018). Mosquito vector-associated microbiota: Metabarcoding bacteria and eukaryotic symbionts across habitat types in Thailand endemic for dengue and other arthropod-borne diseases. Ecology and Evolution, 8(2), 1352–1368. 10.1002/ece3.3676

Tikhonov, G., Abrego, N., Dunson, D., & Ovaskainen, O. (2017). Using joint species distribution models for evaluating how species-to-species associations depend on the environmental context. Methods in Ecology and Evolution, 8(4), 443–452. 10.1111/2041-210x.12723

Toews, D. P. L., & Brelsford, A. (2012). The biogeography of mitochondrial and nuclear discordance in animals. Molecular Ecology, 21(16), 3907–3930. 10.1111/j.1365-294X.2012.05664.x

Tourlousse, D. M., Yoshiike, S., Ohashi, A., Matsukura, S., Noda, N., & Sekiguchi, Y. (2016). Synthetic spike-in standards for high-throughput 16S rRNA gene amplicon sequencing. Nucleic Acids Research, gkw984. 10.1093/nar/gkw984

Venables, W. N., & Ripley, B. D. (2002). Modern Applied Statistics with S. Springer New York. 10.1007/978-0-387-21706-2

Vieira, C. J. S. P., Gyawali, N., Onn, M. B., Shivas, M. A., Shearman, D., Darbro, J. M., Wallau, G. L., Van Den Hurk, A. F., Frentiu, F. D., Skinner, E. B., & Devine, G. J. (2024). Mosquito bloodmeals can be used to determine vertebrate diversity, host preference, and pathogen exposure in humans and wildlife. Scientific Reports, 14(1), 23203. 10.1038/s41598-024-73820-y

Villegas, L. E. M., Radl, J., Dimopoulos, G., & Short, S. M. (2023). Bacterial communities of Aedes aegypti mosquitoes differ between crop and midgut tissues. PLOS Neglected Tropical Diseases, 17(3), e0011218. 10.1371/journal.pntd.0011218

Villeneuve, C.-A., Snyman, L. P., Jenkins, E. J., Lecomte, N., Dusfour, I., & Leighton, P. A. (2024). Variable performance of DNA barcoding and morpholo-gical characteristics for the identification of Arctic black-legged Aedes (Diptera: Culicidae), with a focus on the Punctor subgroup. Arthropod Systematics & Phylogeny, 82, 17–34. 10.3897/asp.82.e111985

Wickham, H., (with Sievert, C.). (2016). ggplot2: Elegant graphics for data analysis (Second edition). Springer.

Williamson, E. M., Hammer, T. J., Hogendoorn, K., & Eisenhofer, R. (2025). Blanking on blanks: Few insect microbiota studies control for contaminants. mBio, 16(4), e02658-24. 10.1128/mbio.02658-24

Yilmaz, P., Parfrey, L. W., Yarza, P., Gerken, J., Pruesse, E., Quast, C., Schweer, T., Peplies, J., Ludwig, W., & Glöckner, F. O. (2014). The SILVA and “All-species Living Tree Project (LTP)” taxonomic frameworks. Nucleic Acids Research, 42(D1), D643–D648. 10.1093/nar/gkt1209

Zhang, J., Kobert, K., Flouri, T., & Stamatakis, A. (2014). PEAR: A fast and accurate Illumina Paired-End reAd mergeR. Bioinformatics, 30(5), 614–620. 10.1093/bioinformatics/btt593

Zhong, J., Jasinskas, A., & Barbour, A. G. (2007). Antibiotic treatment of the tick vector Amblyomma americanum reduced reproductive fitness. PLoS ONE, 2(5), e405. 10.1371/journal.pone.0000405

