## Supplementary material for "Cold and lonely: low microbial abundance and no core microbiota in mosquito populations across Greenland": Suppl. Figures

**Running title:** Greenland mosquito microbiota

Diana Rojas Guerrero^1,2,*^, Tomas Roslin^3,4^, Otso T. Ovaskainen^5,6^, Michał Kolasa^1,7^, Viktor Gårdman^4,8^, Karol H. Nowak^1,2^, Monika Prus-Frankowska^1^, Mateusz Buczek^1^, Anna Michalik^9^, Piotr Łukasik^1,*^

^1^Institute of Environmental Sciences, Faculty of Biology, Jagiellonian University, Kraków, Poland

^2^Doctoral School of Exact and Natural Sciences. PhD Programme in Biology, Faculty of Biology, Jagiellonian University, Kraków, Poland.

^3^Department of Agricultural Sciences, University of Helsinki, Helsinki, Finland

^4^Department of Ecology, Swedish University of Agricultural Sciences, Uppsala, Sweden

^5^Department of Biological and Environmental Science, Faculty of Mathematics and Science, University of Jyväskylä, Jyväskylä, Finland

^6^Organismal and Evolutionary Biology Research Programme, Faculty of Biological and Environmental Sciences, University of Helsinki, Helsinki, Finland

^7^Department of Reproductive Biotechnology and Cryoconservation, National Research Institute of Animal Production, Balice, Poland

^8^Department of Environment and Minerals, Greenland Institute of Natural Resources, Nuuk, Greenland

^9^Institute of Zoology and Biomedical Research, Faculty of Biology, Jagiellonian University, Kraków, Poland

**Correspondence:** Diana Rojas Guerrero, and Piotr Łukasik

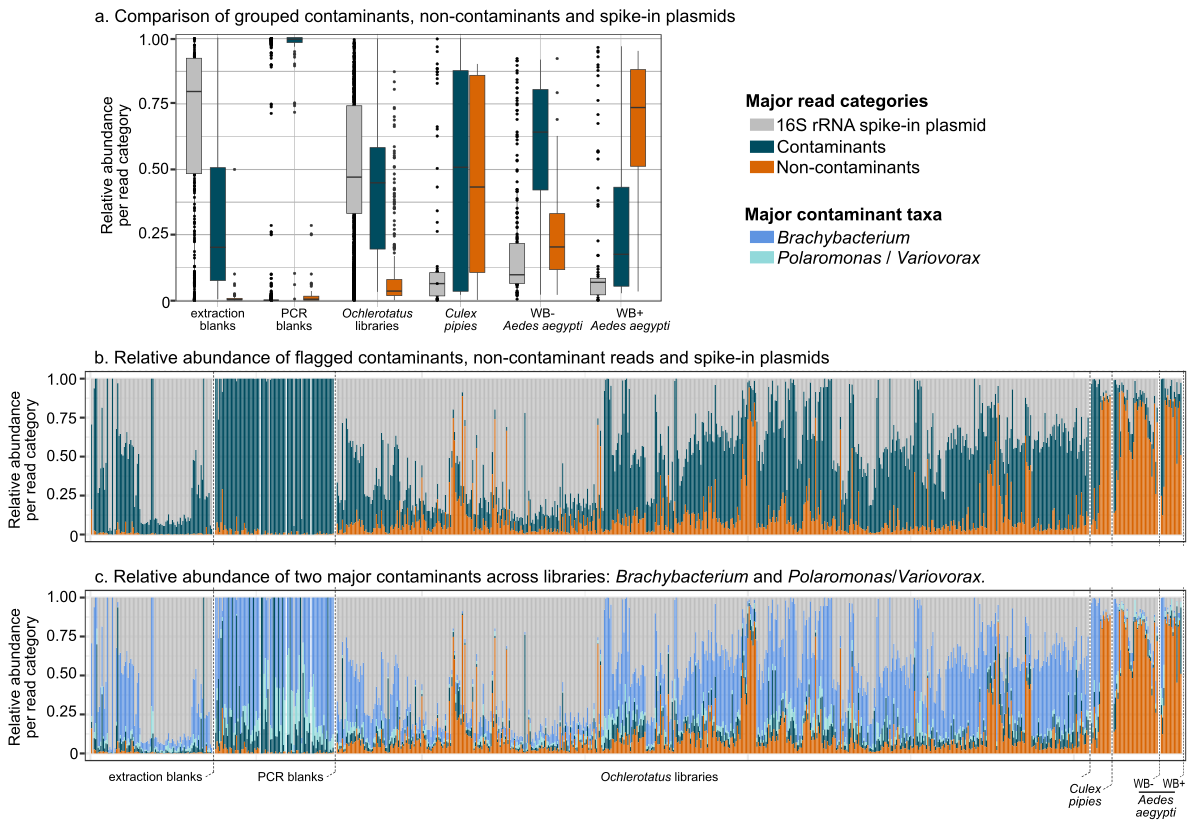

**Fig. S1.** Decontamination of 16S rRNA experimental libraries based on blank controls.

Decontamination was performed on raw libraries (i.e. prior to absolute 16S rRNA copy estimation) as explained in Łukasik et al. (2017). Briefly, after extracting spike-in plasmid reads, raw OTU reads of mosquito libraries and blanks were transformed to relative abundance estimation. Then, the maximum abundance of each OTU was estimated independently for both types of libraries. Maximum abundance values of OTUs in blanks were multiplied by ten and compared against maximum abundance values of OTUs in mosquito libraires. If OTU abundance across blanks was equal or higher than in experimental libraries, then the OTU was classified as “contaminant”, otherwise, it was flagged as a “non-contaminant”.

a) Boxplots showing the median read categories across library types. *Ochlerotatus* libraries showed a high number of spike-ins and contaminants in comparison with the detected non-contaminant reads, resembling extraction blanks. On the other hand, other mosquito species showed a high bacterial load, regardless of whether *Wolbachia* was present (i.e. *C. pipiens*, WB+ *A. aegypti*) or not (WB- *A. aegypti*). In contrast, the spike-in detection in such libraries was lower, in comparison with blanks and *Ocheloratus* libraries.

b) Relative abundance of spike-ins, contaminants, and non-contaminants per library before decontamination revealed a high proportion of contaminants in *Ochlerotatus* libraries.

c) Relative abundance of the two dominant contaminants. *Brachybacterium*, often found in PCR reagents (refs), and an OTU classified as *Polaromonas*, which included ASVs resembling *Variovorax* sp. - a taxon linked to mosquito microbiota, were also present in blank libraries and thus flagged as contaminants.

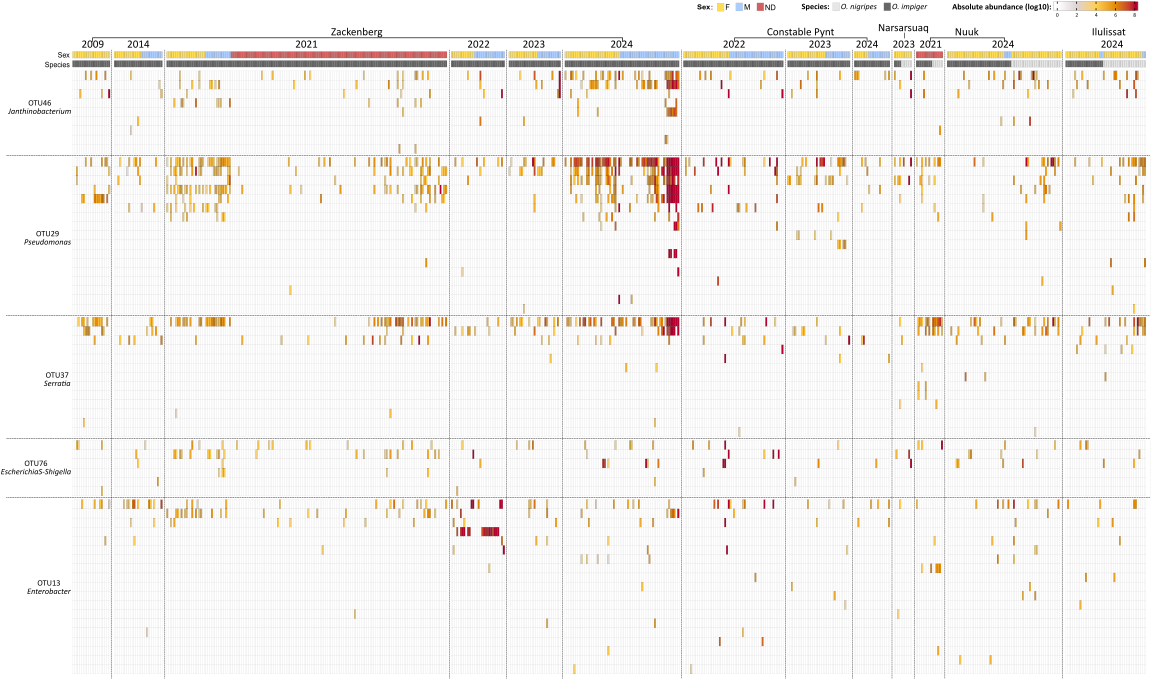

Kobbefjord

**Fig. S2**. Bacterial genotype diversity for the five most abundant and prevalent taxa across mosquito populations from multiple locations and years. Estimated absolute abundances are shown at the OTU-clustered level with taxonomic classifications for ASVs detected in more than one library, based on 573 mosquito individuals.

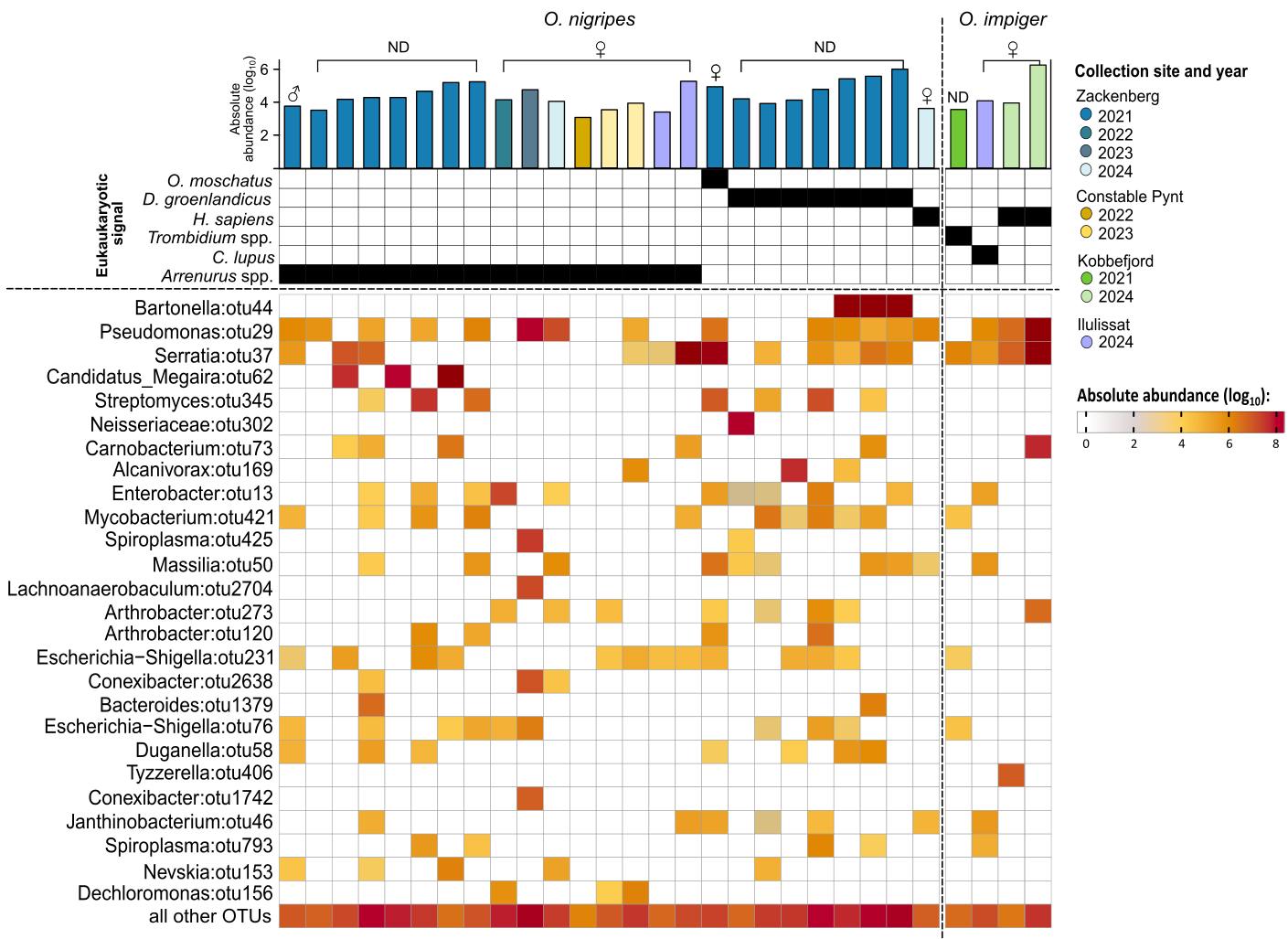

**Fig. S3.** Overview of mosquito libraries containing non-mosquito eukaryotic signal and their associated bacterial communities*. From top to bottom: Bar plots showing total bacterial load (estimated 16S rRNA gene copy number) for each individual mosquito, with bar colors indicating collection site; tile plot showing the presence of non-mosquito eukaryotic sequences detected in each library; and heatmap displaying absolute 16S rRNA gene copy numbers for the most abundant bacterial OTUs across the mosquito libraries carrying eukaryotic signal.

*****Non-*Ochlerotatus* COI reads (0.41% of the dataset) included: water mites *Arrenurus* (96,208 reads, n = 16), and *Trombidium* mites (1,985 reads, n = 1), Northern collared lemming *Dicrostonyx groenlandicus* (9,615 reads, n = 7), muskox *Ovibos moschatus* (27 reads, n = 3), Greenland dog *Canis lupus* (998 reads, n = 1), and human *Homo sapiens* (5,365 reads, n = 3). Human COI reads in the affected libraries shown here had >700 *H. sapiens* reads per library. 36 further libraries contained human COI reads at much lower abundance (2-12 reads), but we conservatively assumed that they may represent low-level background contamination and did not include them here.

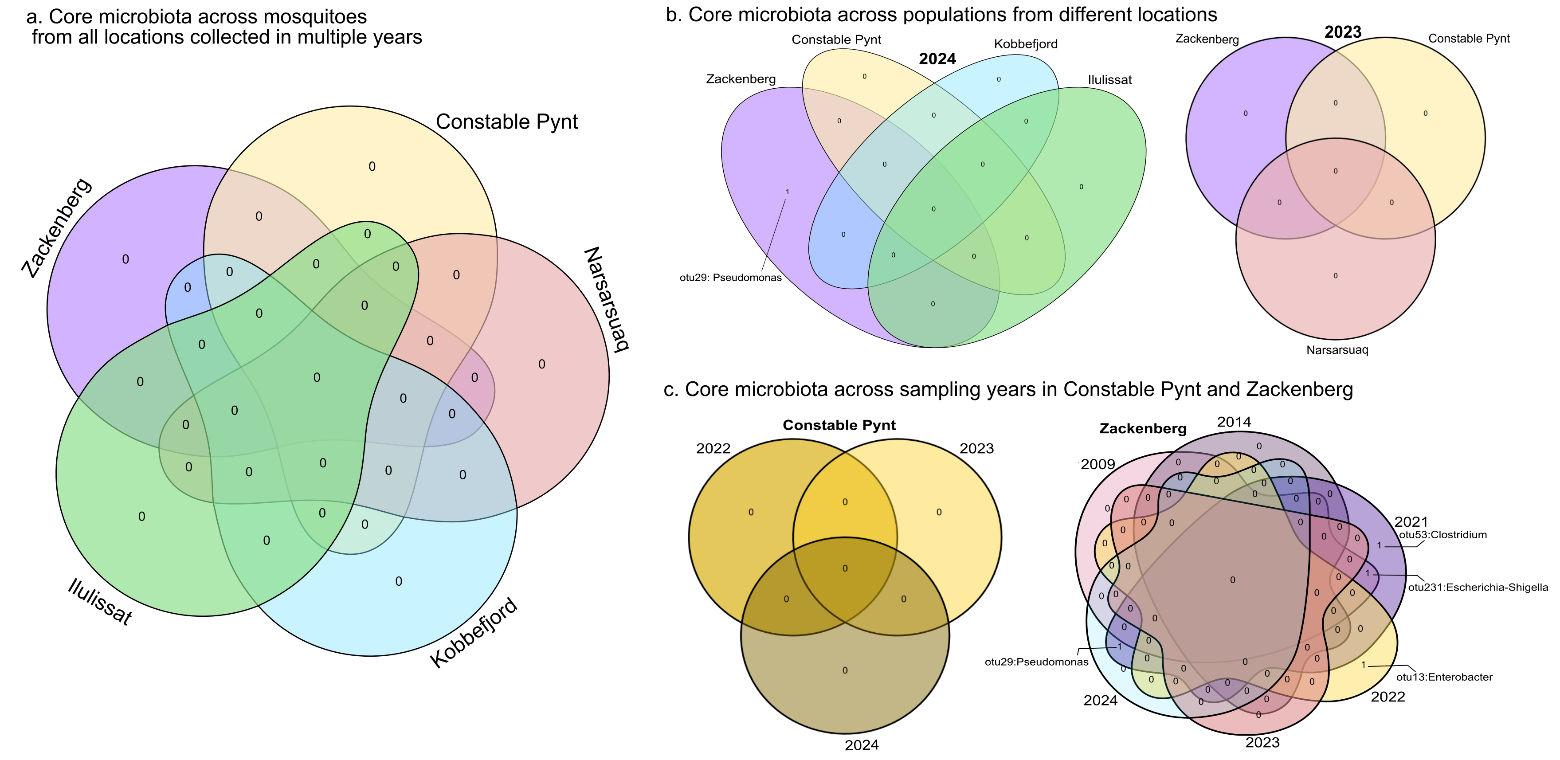

**Fig. S4.** Core microbiota across mosquito populations sampled in different years and locations, detected by 16S rRNA amplicons, using a threshold of 0.01% bacterial abundance at 75% prevalence. **a)** Across locations (regardless of year), no taxa met the core criteria. **b)** When considering year and location together, *Pseudomonas* was the only prevalent taxon, detected in Zackenberg 2024. **c)** Temporal analysis at Constable Pynt and Zackenberg revealed some taxa shared between years, including *Pseudomonas* and *Escherichia–Shigella*.

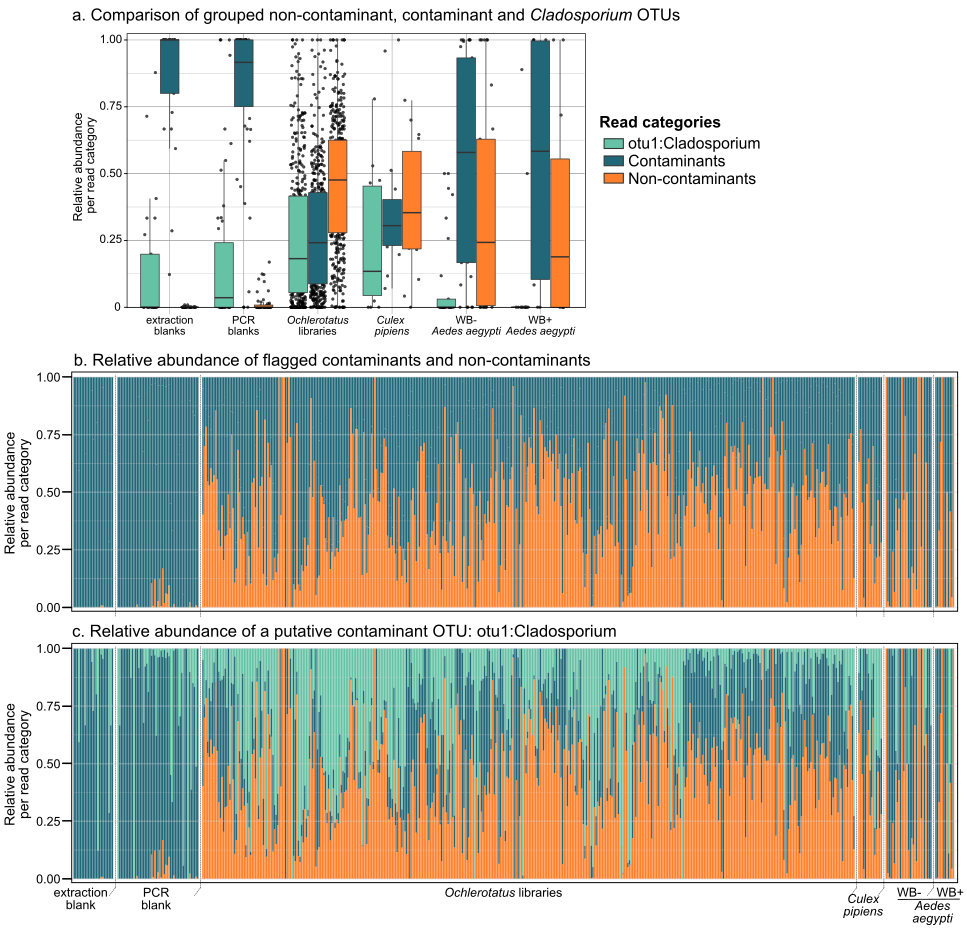

**Fig. S5.** Decontamination of fungal ITS2 experimental libraries based on blank controls.
Decontamination was carried out as stated for 16S rRNA amplicons, and described in detail in (Łukasik *et al.*, 2017).

a) Boxplots showing the proportion of read categories across library types.Beause Cladosporium was the most abundant OTU, equally present across libraries, it was assigned as an independent category. *Ochlerotatus* libraries had the highest amount of non-contaminant reads making up almost 50% all the different categories. At the same time, and in comparison with the other mosquito species libraires, Ochlerotatus libraries had also the highest *Cladosporium* number of reads.
b) Relative abundance of all contaminants and non-contaminants across library types.
c) Relative abundance of Cladosporium across library types.

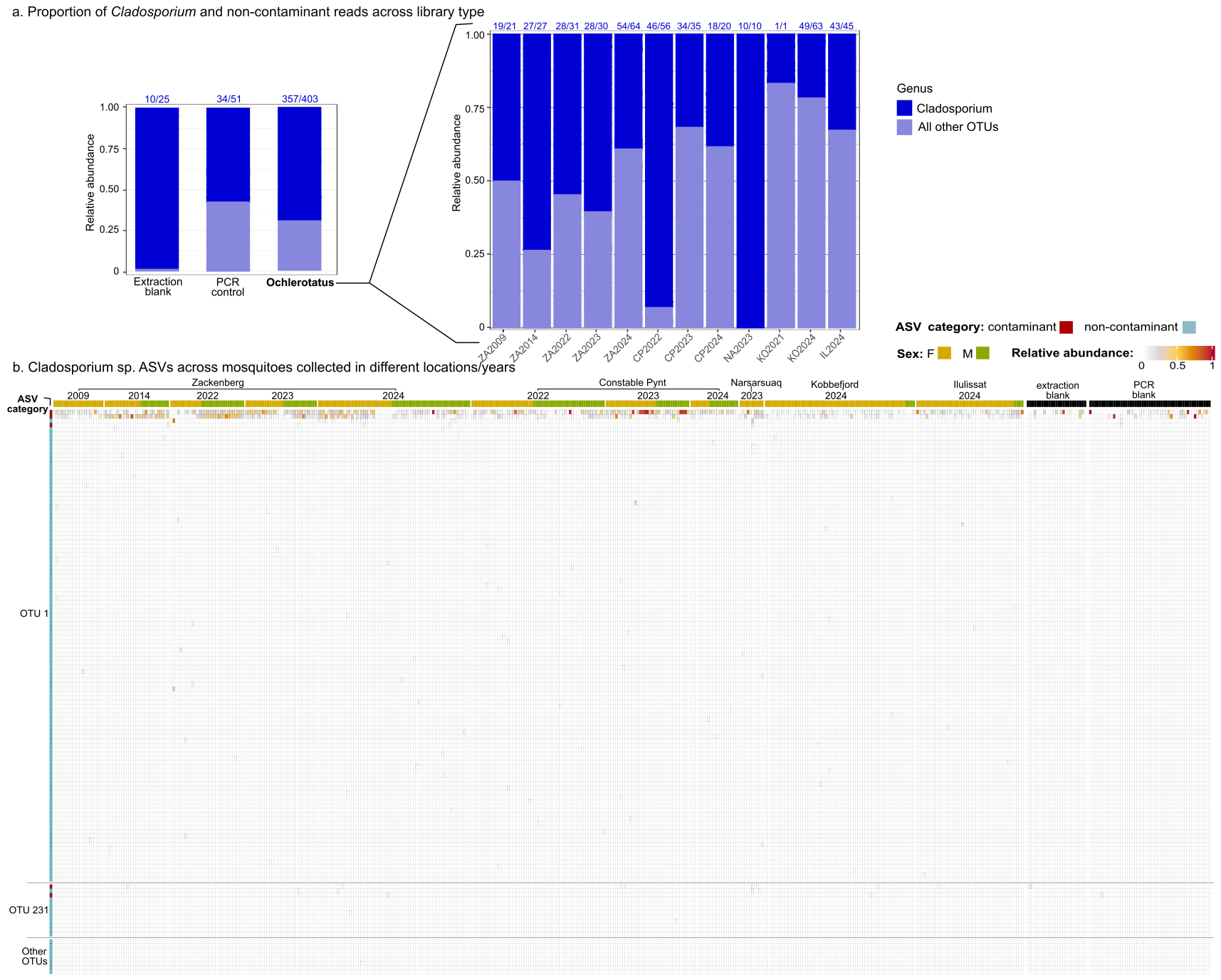

**Fig. S6.** *Cladosporium* prevalence and abundance across library types. a) Summary of abundance and frequency of *Cladosporium* OTUs across control and experimental libraries. The numbers on top of the bars indicate the number of libraries where at least one *Cladosporium* read was detected/total number of libraries. b) Relative abundance of *Cladosporium* ASVs across individual libraries and library types. Rows represent an individual *Cladosporium* genotype (ASV) clustered into specific OTUs. The first column indicates whether, after the decontamination process, the individual ASV was flagged as a contaminant or not. Subsequent columns represent individual libraries and relative abundances for individual ASVs. The ‘Other OTUs’ category encompasses individual OTUs with no ASVs.

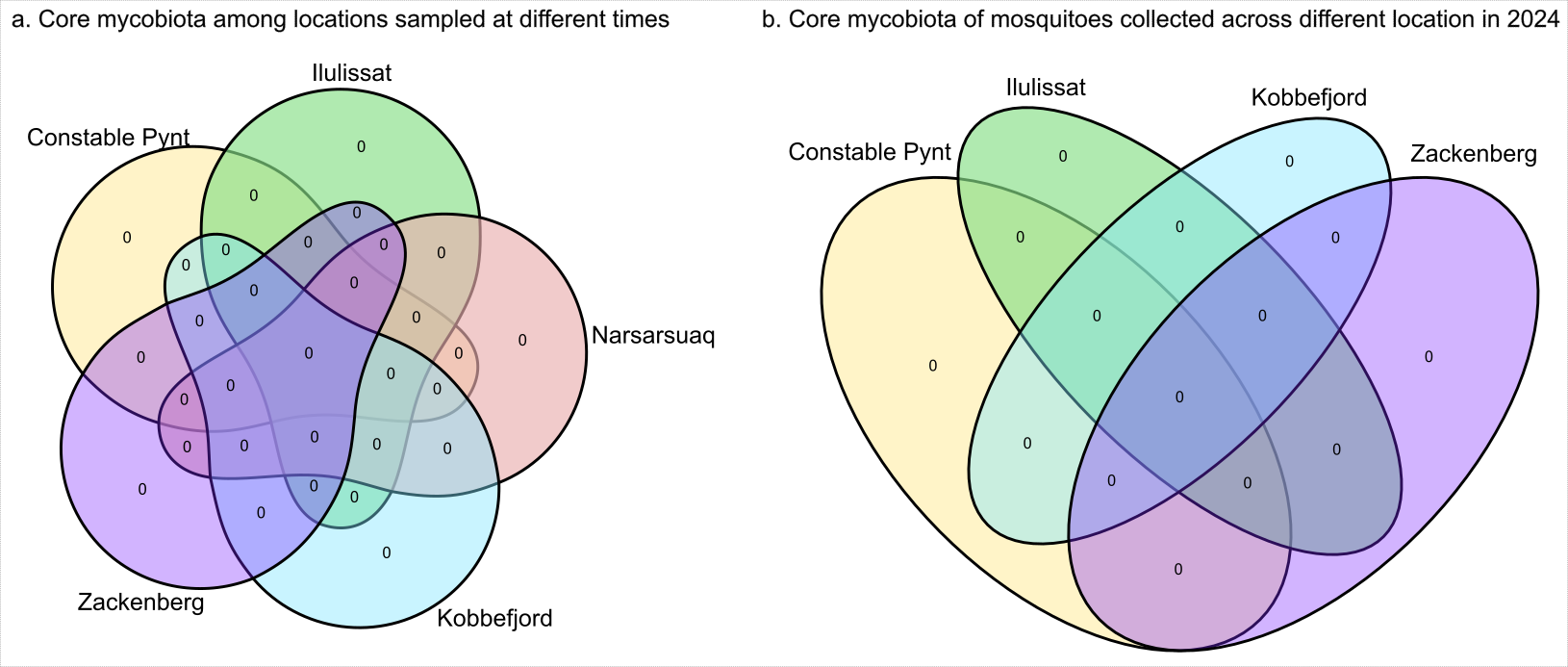

**Fig. S7.** Core microbiota across mosquito populations sampled in different years and locations, detected by ITS2 amplicons, using a threshold of 0.01% fungal abundance at 75% prevalence. a) Across locations (regardless of year), no taxa met the core criteria. b) Core microbiota of mosquitoes sampled across Greenland in 204.

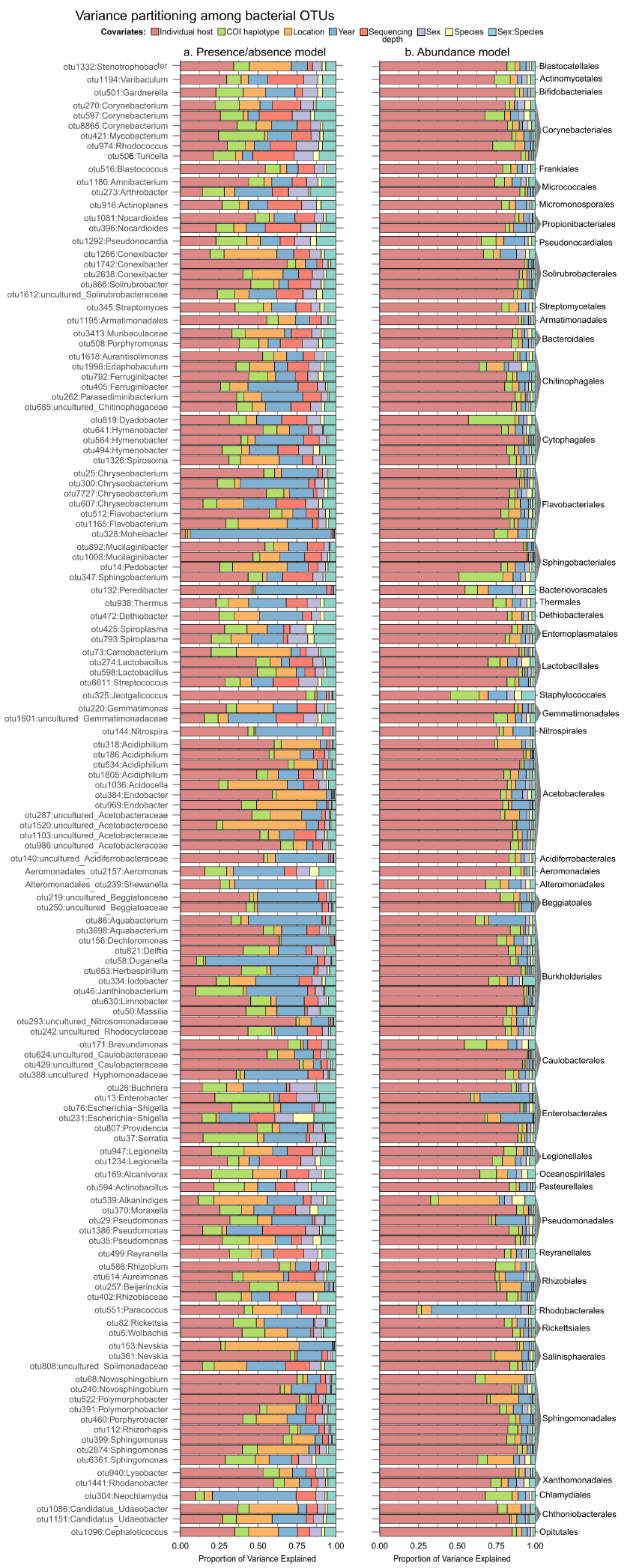

**Fig. S8.** Variance partitioning among the fixed and random effects included in two HMSC models for the top 100 most abundant bacterial OTUs. Each bar represents an individual 16S OTU and the proportion of explained variance attributed to random (i.e. host individual, COI haplotype, location, year) or fixed (i.e. sex, species, or their interaction) effects. Panel show variance partitioning for the a) presence/absence model and b) abundance conditional on presence model. Variance partitioning for all bacterial OTUs is available in Table S5.

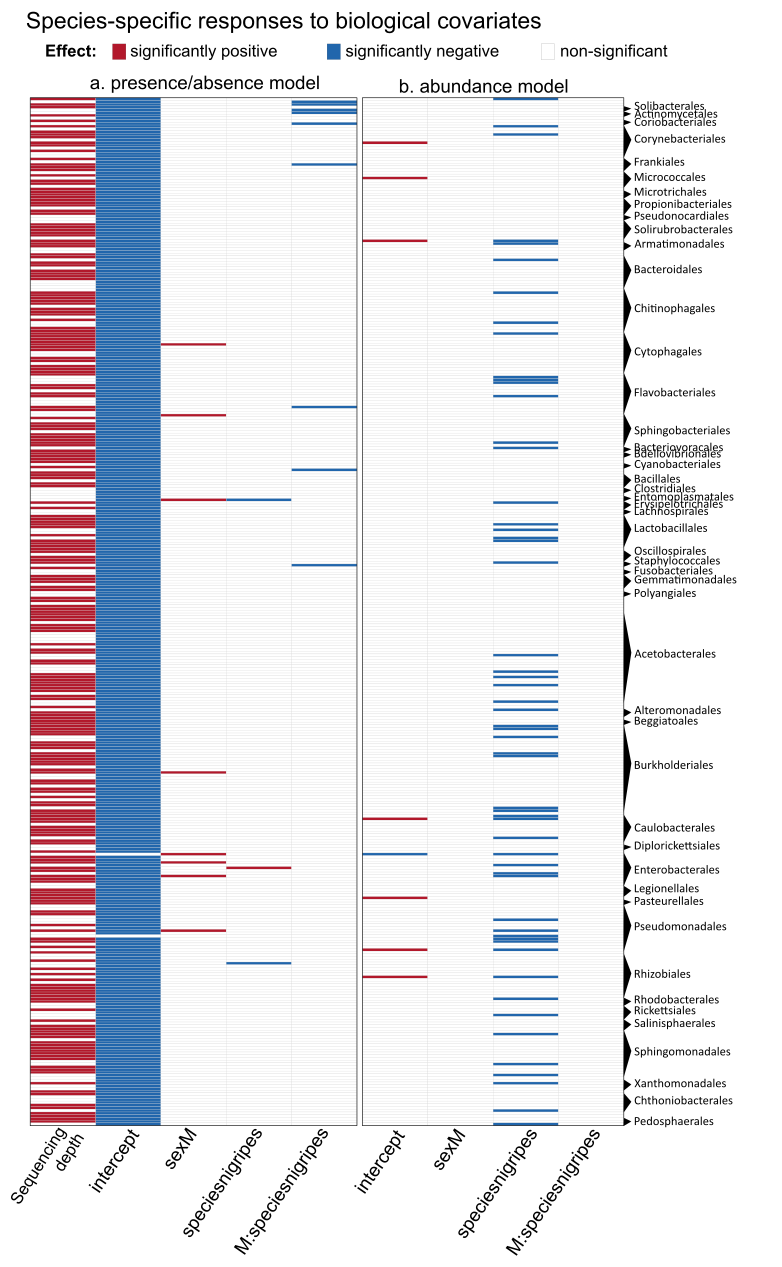

**Fig. S9.** Species-specific responses to biological covariates across all sampled bacterial OTUs. The columns show the posterior probability that the effect in question deviates from zero, with red tiles indicating significantly positive effects, blue tiles indicating significantly negative effects, and white tiles indicating non-statistically significant responses (posterior probability > 0.95). Note that, given the model's parameterisation, tests relate to the difference between the factor in question and the reference group (i.e., females of *O. impiger*). Thus, the colours refer to differences between this group and males of *O. impiger;* females of *O. nigripes*, and males *O. nigripes*. The panels show results for a) the presence/absence model, and b) for the model of abundance conditional on presence. Full taxonomic annotation for all bacterial OTUs available in Table S6.

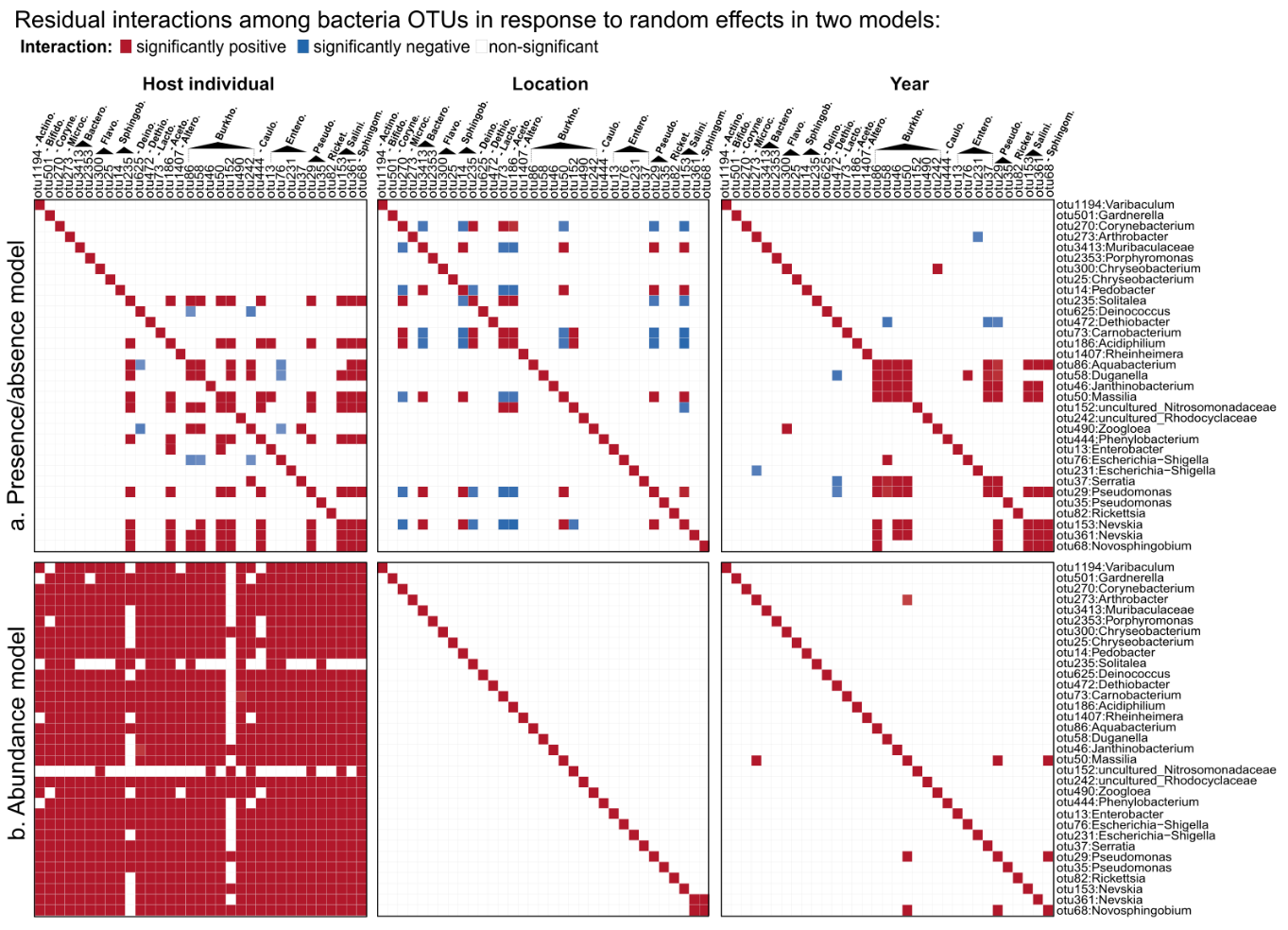

**Fig. S10.** Residual species-to-species associations across the top 33 most prevalent bacterial OTUs. Each panel shows the association patterns among bacterial OTUs, at different levels in two HMSC models: a) presence/absence and b) abundance conditional on presence. Red indicates a significant positive association and blue a significant negative association (posterior probability > 0.95). Full dataset interactions are available in at <https://figshare.com/s/fae30efc070ea5e7535c>.

Considering that interactions could be a result of taxa responding similarly to shared environmental or host-related factors, and from direct or indirect interactions among taxa themselves, our HMSC models first accounted for joint responses to extrinsic factors (fixed and random effects). After controlling for these effects, we examined the residual species-species associations that cannot be attributed to the measured covariates. These associations were examined at multiple hierarchical levels: host individual, host haplotype, location, and year.

In terms of species occurrence, relatively few taxa showed statistically supported associations, but within the 30 most abundant and prevalent OTUs, several statistically supported interactions emerged (Fig. S10a). At the host-individual level, interactions were mostly positive, except for *Escherichia-Shigella*, which tended to co-occur negatively with environmentally associated taxa like *Zooglea, Aquabacterium*, and *Duganella*. At the location level, most of the statistically supported interactions were negative; for example, *Pseudomonas* and *Massilia* were negatively associated with *Carnobacterium*, despite all three being present across populations. At the year level, members of the prevalent set (e.g., Escherichia-Shigella, *Janthinobacterium*, *Massilia, Pseudomonas*) showed strong positive associations. No statistically supported associations were observed at the haplotype level.

In terms of abundance conditional on presence, statistically supported residual associations were positive but rare at all levels, except at the host individual level (Fig. S10b). Notably though, most members of the prevalent set of microbial taxa showed positive residual associations with each other across different levels in the occurrence model (Fig. S10b). At the location level, only *Novosphingobium* and a specific *Nevskia* genotype showed a positive association. At the year level, *Massilia* showed a residual association with *Pseudomonas* and *Novosphingobium* in terms of both occurrence and abundance conditional on presence. However, at the host-individual level, nearly all taxa showed positive associations in terms of abundance. Thus, if an individual mosquito showed high abundances of one microbe, it tended to show high abundances of other taxa, too. At the level of mosquito haplotypes, no significant associations were detected for either the presence/absence or abundance conditional on presence of microbial taxa (Fig. S10b).

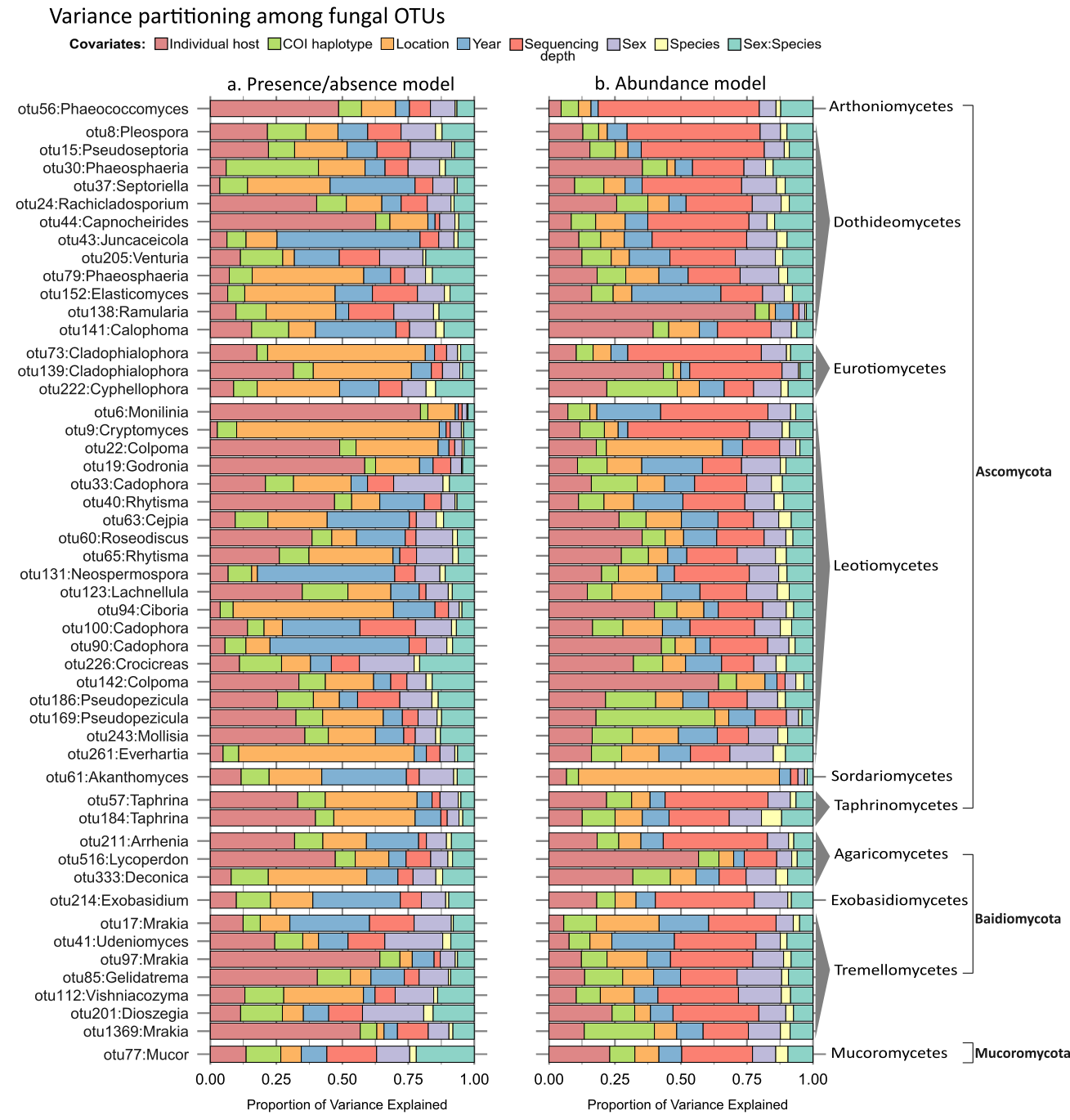

**Fig. S11.** Variance partitioning among the fixed and random effects included in two HMSC models. Each bar represents an individual ITS2 OTU and its variance proportions as explained by random (i.e. host individual, COI haplotype, location, year) or fixed (i.e. sex, species, or their interaction) effects. Panel show variance partitioning for the a) presence/absence model and b) abundance conditional on presence model.

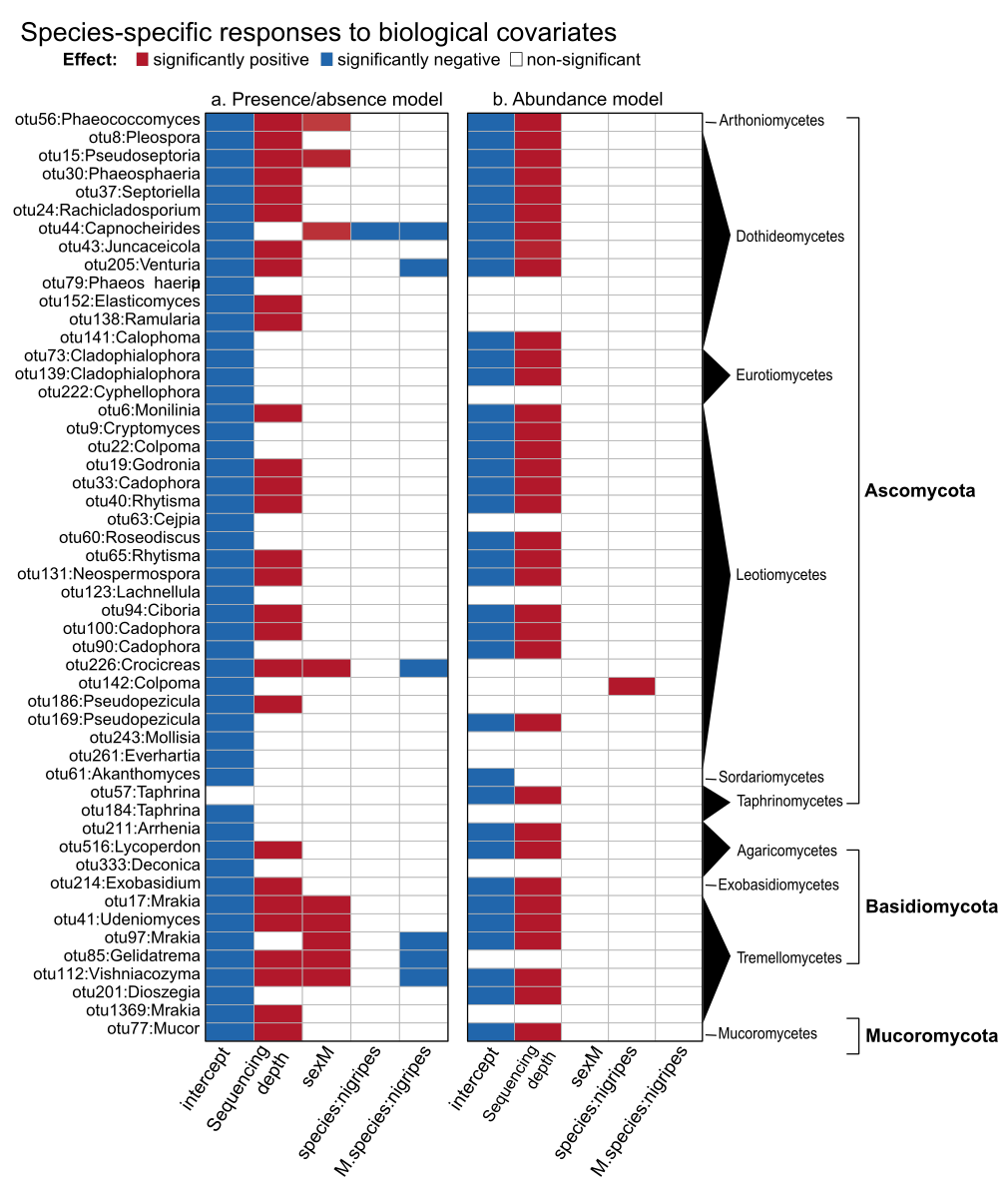

**Fig. S12.** Species-specific responses to biological covariates across all sampled fungal OTUs. The columns show the posterior probability that the effect in question deviates from zero, with red tiles indicating significantly positive effects, blue tiles indicating significantly negative effects, and white tiles indicating non-statistically significant responses (posterior probability > 0.95). Note that, given the model's parameterisation, tests relate to the difference between the factor in question and the reference group (i.e., females of *O. impiger*). Thus, the colours refer to differences between this group and males of *O. impiger;* females of *O. nigripes*, and males *O. nigripes*. The panels show results for a) the presence/absence model, and b) for the model of abundance conditional on presence. Full taxa annotation in Table S6.

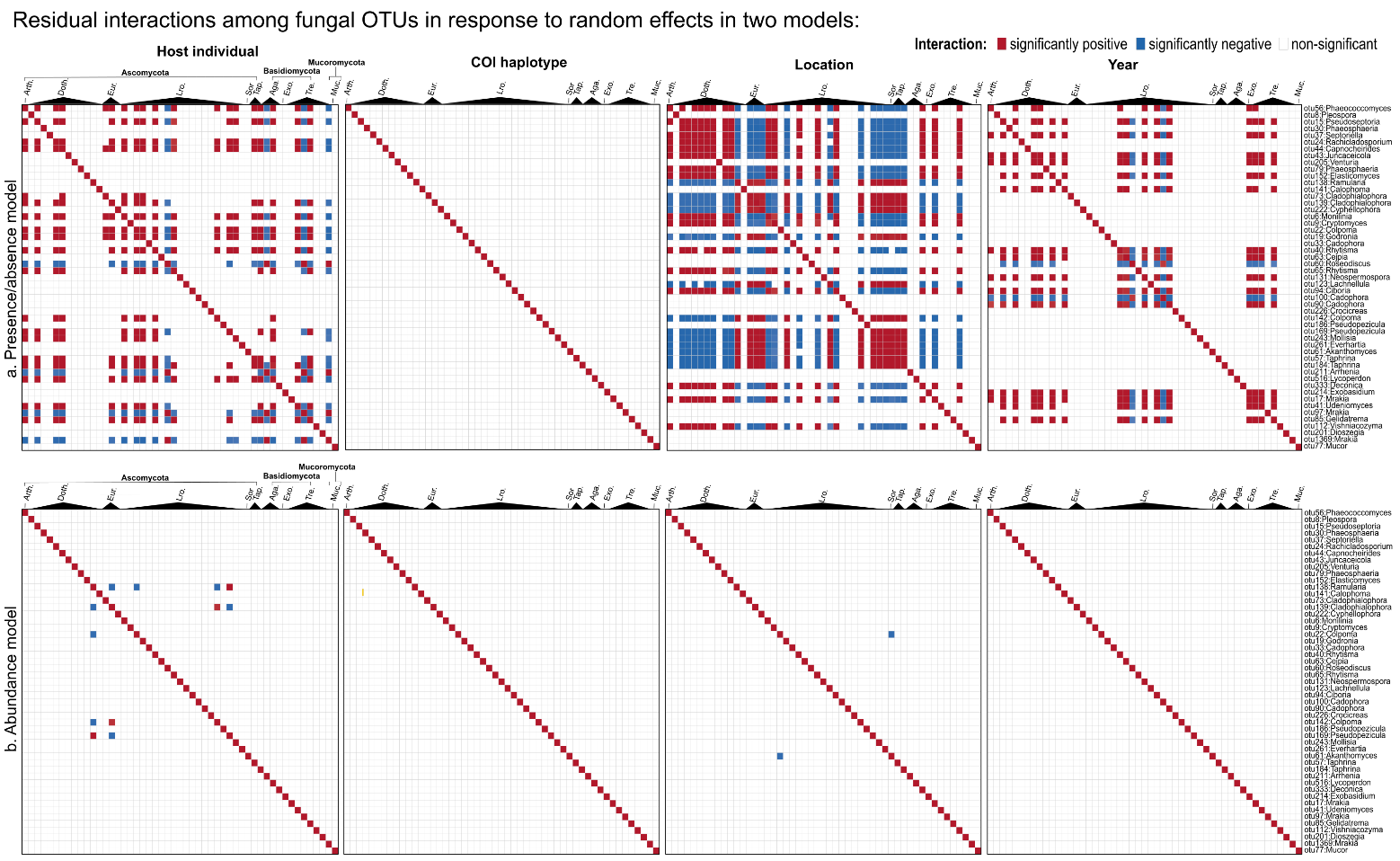

**Fig. S13.** Residual species-to-species associations. Included are all fungal species. Each panel shows the association patterns among bacterial OTUs, at different levels in two HMSC models: a) presence/absence and b) abundance conditional on presence. Red indicates a significant positive association and blue a significant negative association (posterior probability > 0.95).

Several fungal taxa showed statistically supported residual associations across all the 51 analysed OTUs (Fig. S13) in the occurrence model. Following the most prevalent OTUs across mosquitoes (e.g*., Mrakia, Monilinia, Cryptomyces* and, *Phaeosphaeria*; see Fig. 5c), most detected associations were positive at the level of the host individual. Monilinia appeared to interact positively with site-specific taxa such as *Colpoma* and *Godronia*, while these three taxa showed negative associations with other environmental fungi like *Poseodiscus* and Arrhenia (Fig. S13a). At the location level, interactions were more mixed, with *Monilinia* and *Mrakia* positively associated with each other and with *Cryptomyces,* but negatively or not associated with *Colpoma* and *Godronia*. Notably, none of these taxa displayed statistically supported interactions at the year level. At the haplotype level, we found no associations at all.

In terms of abundance conditional on presence, statistically supported residual associations tended to be scarce at any level (Fig. S13b).

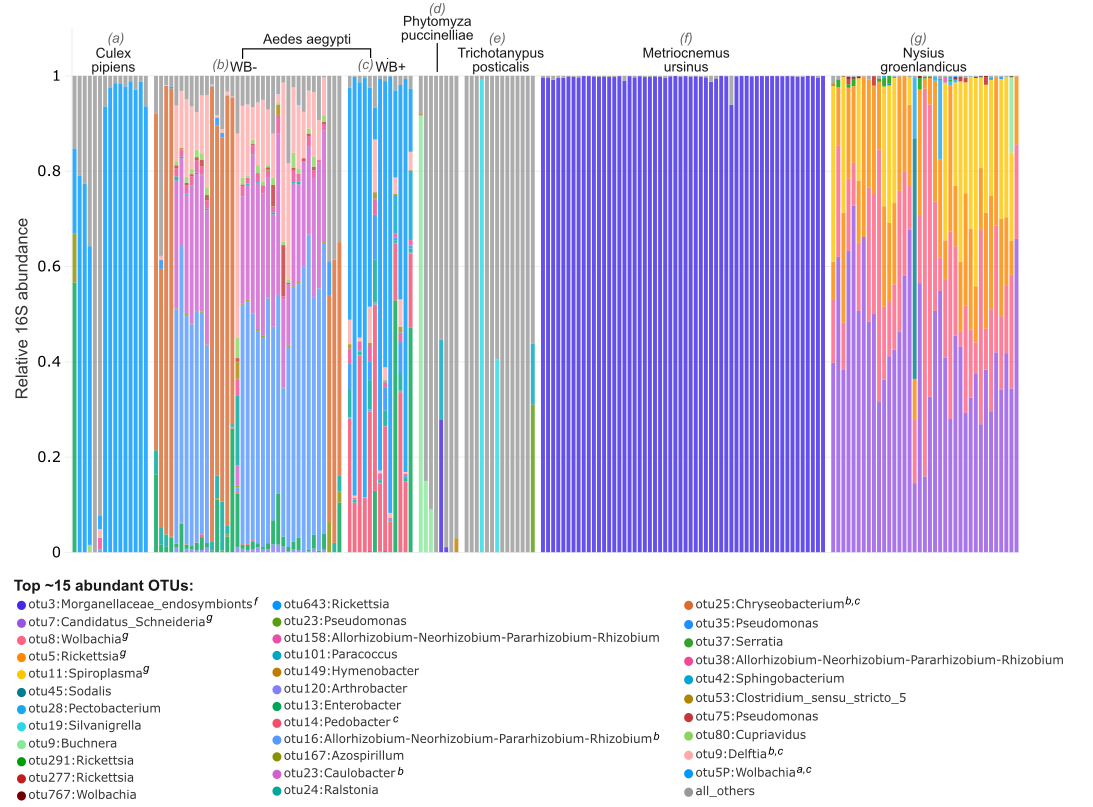

**Fig. S14.** Relative abundance of bacterial OTUs in reference species, including wild *C. pipiens*, lab cultured *Aedes aegypti* with or without *Wolbachia* infection. Additionally, we included dipteran species *P. puccinelliae*, *T. posticalis*, and *M. ursinos*, as well as the hemipteran *N. groenlandicus*, collected in Zackenberg in 2021. OTUs displayed were classified at the genus level and include the top ~15 most abundant OTUs across mosquito species and Greenlandic species.
